# Programmable RNA detection with CRISPR-Cas12a

**DOI:** 10.1101/2023.01.29.525716

**Authors:** Santosh R. Rananaware, Emma K. Vesco, Grace M. Shoemaker, Swapnil S. Anekar, Luke Samuel W. Sandoval, Katelyn S. Meister, Nicolas C. Macaluso, Long T. Nguyen, Piyush K. Jain

## Abstract

CRISPR is a prominent bioengineering tool and the type V CRISPR-associated protein complex, Cas12a, is widely used in diagnostic platforms due to its innate ability to cleave DNA substrates. Here we demonstrate that Cas12a can also be programmed to directly detect RNA substrates without the need for reverse transcription or strand displacement. We discovered that while the PAM-proximal “seed” region of the crRNA exclusively recognizes DNA for initiating *trans-*cleavage, the PAM-distal region or 3’-end of the crRNA can tolerate both RNA and DNA substrates. Utilizing this property, we developed a method named Split Activators for Highly Accessible RNA Analysis or ‘SAHARA’ to detect RNA sequences at the PAM-distal region of the crRNA by merely supplying a short ssDNA or a PAM containing dsDNA to the seed region. Notably, SAHARA is Mg^2+^ concentration- and pH-dependent, and it was observed to work robustly at room temperature with multiple orthologs of Cas12a. SAHARA also displayed a significant improvement in the specificity for target recognition as compared to the wild-type CRISPR-Cas12a, at certain positions along the crRNA. By employing SAHARA we achieved amplification-free detection of picomolar concentrations of miRNA-155 and hepatitis C virus RNA. Finally, SAHARA can use a PAM-proximal DNA as a switch to control the trans-cleavage activity of Cas12a for the detection of both DNA and RNA targets. With this, multicomplex arrays can be made to detect distinct DNA and RNA targets with pooled crRNA/Cas12a complexes. In conclusion, SAHARA is a simple, yet powerful nucleic acid detection platform based on Cas12a that can be applied in a multiplexed fashion and potentially be expanded to other CRISPR-Cas enzymes.

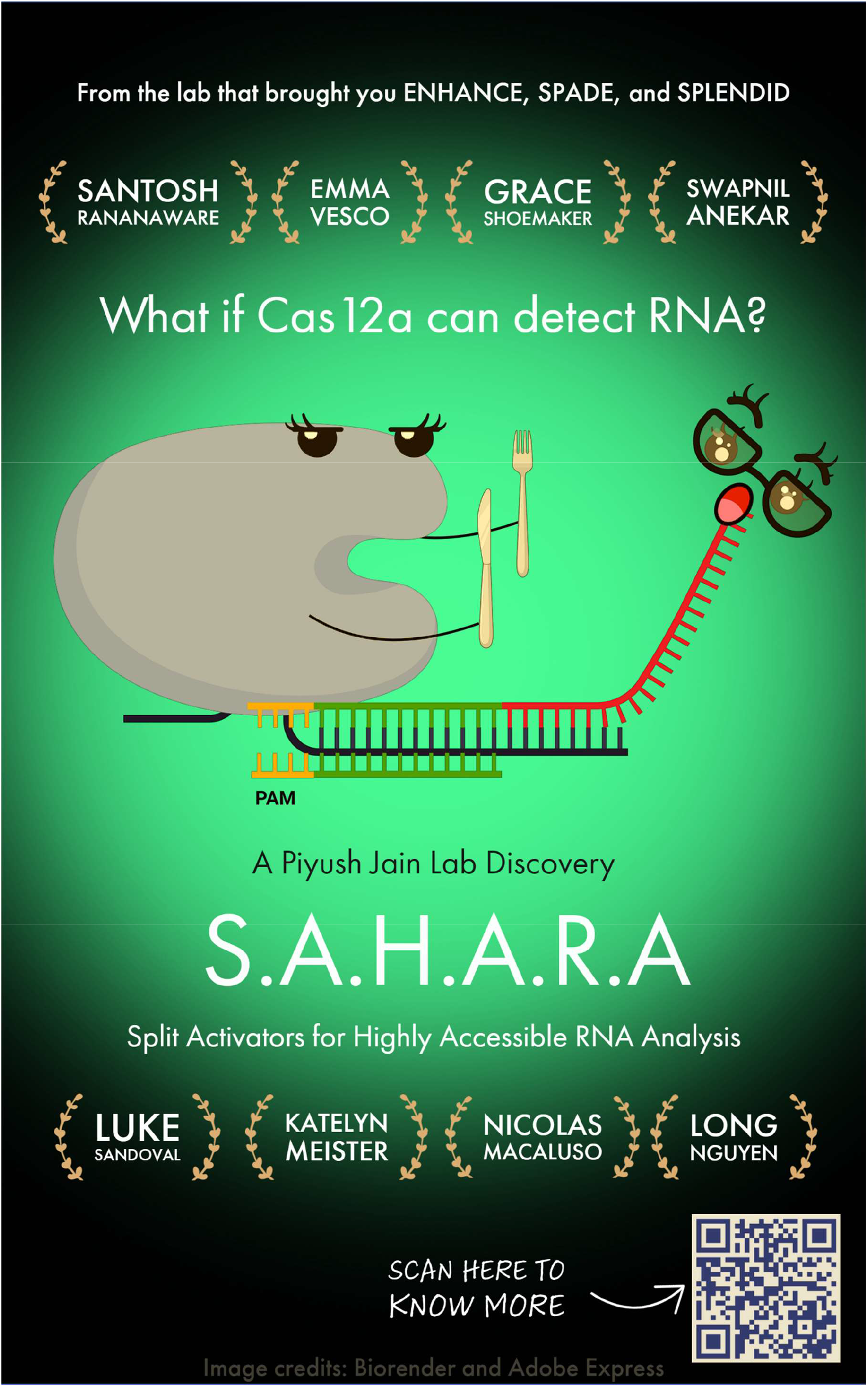

## Introduction

CRISPR (Clustered Regularly Interspaced Short Palindromic Repeats) is an adaptive immune system encoded within prokaryotes that have evolved to counter invasion by foreign nucleic acids such as bacteriophages and plasmids^1,2^. Upon infection, the invading DNA sequences are captured and integrated into the host genome between an array of repeat sequences. The captured DNA sequences are called ‘spacers’ and they provide a genetic memory of prior infections^3^. For prokaryotic immunity, the CRISPR locus is transcribed and processed to generate multiple mature CRISPR RNAs (crRNA), each encoding a unique spacer sequence. Cas (CRISPR-associated) proteins are RNA-guided endonucleases that when complexed with the CRISPR RNAs (crRNAs) can enable the cleavage of nucleic acids that are complementary to the crRNA sequence.

There are several diverse naturally occurring CRISPR/Cas systems found in prokaryotes^4,5^. Among these, Cas12a is a class II, type V RNA-guided DNA endonuclease^6^. Since its discovery, it has been widely used for genome editing as well as molecular diagnostic applications^7–11^. Structural and biochemical studies have shown that Cas12a can catalyze the cleavage of DNA substrates^12–14^ but there are no reports of targeted RNA cleavage by Cas12a. Recently there has been a report of a novel enzyme named Cas12a2 that sometimes co-occurs with Cas12a systems in bacteria and can use the Cas12a crRNA, but recognizes an RNA target instead of a double-stranded DNA^15^. A special characteristic of the type V Cas12 family of enzymes is their ability to initiate rapid and indiscriminate cleavage of any non-specific single-stranded DNA (ssDNA) molecules in their vicinity after target-specific recognition and cleavage^16,17^. This unique catalytic property, known as *trans-*cleavage has been harnessed to engineer CRISPR-based diagnostic tools that rely on the cleavage of FRET reporters upon target recognition^18^.

So far, Cas12a-based tools have been limited to the detection of DNA substrates, unless they are coupled with additional steps involving reverse transcription or strand displacement^19–21^. Recently, we discovered that CRISPR-Cas12a can tolerate DNA/RNA heteroduplexes, but only when RNA is located at the non-target strand^22^. We and others utilized the system to detect RNA targets with Cas12a by simply creating a heteroduplex using a reverse transcription step without amplification^22,23^. A reverse-transcription step is inconvenient because it adds to the time, cost, error, and complexity of the assay. The other alternative is to use an RNA-targeting enzyme such as Cas12a2^15^, Cas12g^24^, or Cas13a-d^25–27^; however, these systems can only detect RNA.

Cas12a orthologs are known to require a short protospacer adjacent motif (PAM) to be present on the target DNA to initiate recognition and cleavage^6^. It has been previously shown that DNA-cleaving enzymes such as Cas9 can be manipulated to also cleave RNA through the addition of a PAMmer sequence^28^. However, similar approaches have not yet been investigated with *trans-*cleaving Cas enzymes like Cas12, primarily because the PAM recognition mechanism is very different between Cas9 and Cas12 enzymes^29^. For instance, unlike Cas9, Cas12a can recognize and even trigger *trans-*cleavage activity using a PAM-less ssDNA. To date, there is no single CRISPR-Cas system identified that can innately tolerate both DNA and RNA substrates to trigger *trans-*cleavage.

In this report, we discovered that Cas12a can also tolerate RNA substrates at the PAM-distal end of the crRNA and initiate *trans-*cleavage activity. In essence, we have found that while the PAM-proximal seed region of the crRNA strictly tolerates DNA substrates, the PAM-distal end of the crRNA can tolerate both RNA and DNA substrates with multiple Cas12a orthologs. Thus, by merely supplying a short ssDNA or a PAM-containing dsDNA at the seed region of the crRNA we can detect RNA substrates at the 3’-end of the crRNA. We harnessed this unique property to develop a tool for RNA detection using Cas12a named <u>S</u>plit <u>A</u>ctivators for <u>H</u>ighly <u>A</u>ccessible <u>R</u>NA <u>A</u>nalysis (SAHARA).

We achieved reverse transcription (RTx)-free detection of picomolar levels of DNA as well as RNA without amplification using SAHARA and applied it for detecting clinically-relevant targets including hepatitis C virus (HCV) RNA and microRNA-155 (miR-155). We showed that compared to conventional CRISPR-Cas12a, SAHARA has improved specificity and can be performed at room temperature. We demonstrate that its activity can be turned ON or OFF using the seed region binding DNA activator as a switch. We took advantage of this switch to perform multiplexed and simultaneous detection of different DNA and RNA targets. We also coupled SAHARA with Cas13b to perform multiplexed detection of different RNA targets. These key findings provided insights into the substrate requirements for the *trans-*cleavage activity of Cas12a, and we have utilized them to develop SAHARA, a valuable and versatile tool that can simultaneously detect both DNA and RNA substrates.

## Results

### Cas12a orthologs tolerate split ssDNA activators for *trans-*cleavage activity

LbCas12a, AsCas12a, and ErCas12a are orthologs of Cas12a nucleases that are derived from the *Lachnospiraceae bacterium* ND2006, *Acidaminococcus sp*. BVL36, and *Eubacterium rectale* are simply referred to here as Lb, As, and Er, respectively^30–33^ (Fig. S1). The mature crRNAs for each ortholog are 41-44 nt in length, containing ~19-21 nt of scaffold and the remaining ~20-24 nt of spacer^6^.

We wondered whether two different ssDNA targets, each binding to a different position of the same crRNA, can be used to initiate the *trans-*cleavage activity of Cas12a (Fig. 1a, S2). To test the minimum length of activators tolerated by Cas12a, we designed several short ssDNA target activators of lengths ranging from 6-20 nt that were complementary to either the PAM-proximal (Pp) seed region or the PAM-distal (Pd) end of the crRNA (Fig. 1b-d).

**Fig. 1:**
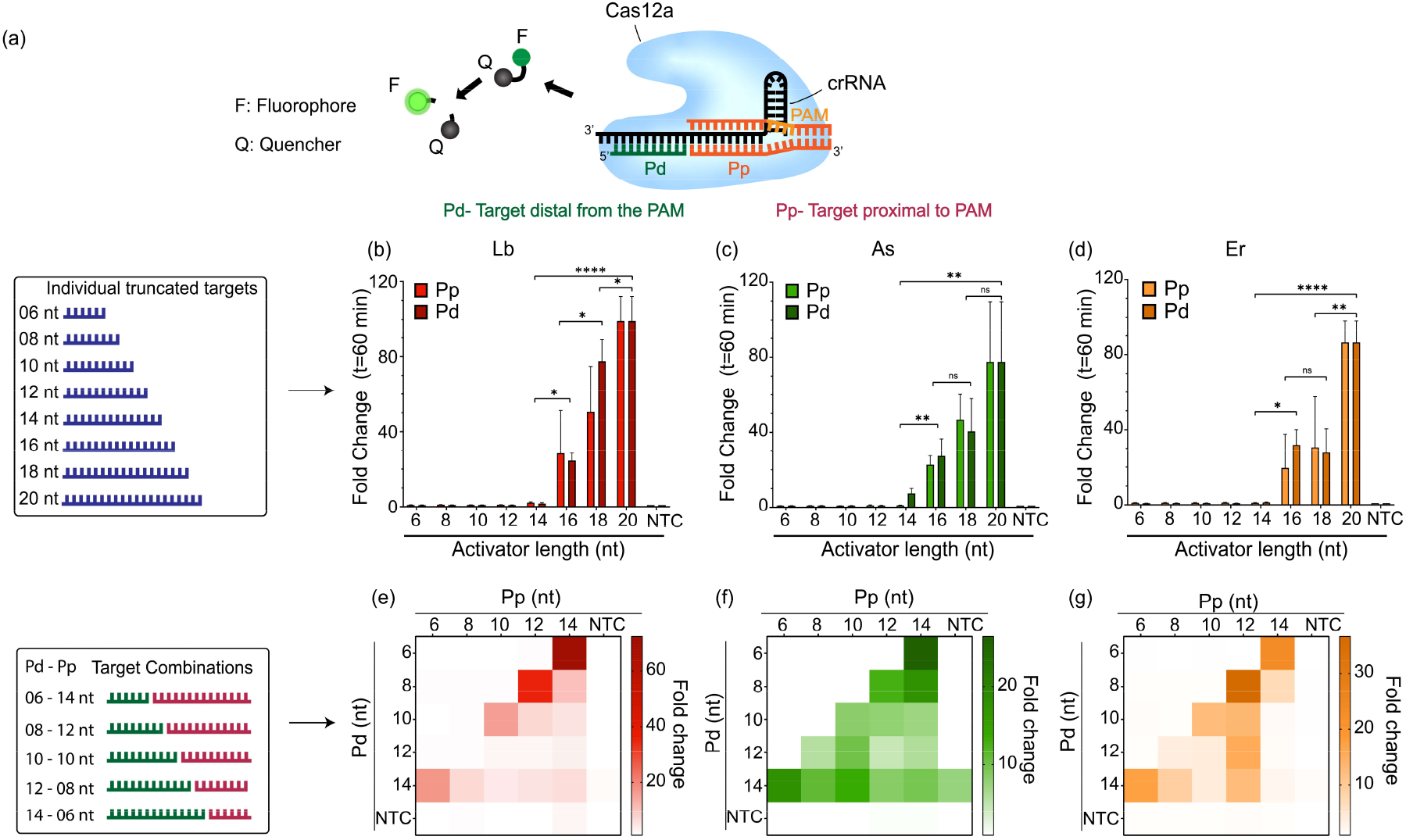
Cas12a orthologs tolerate short ssDNA activators (6-12 nt) when added in combination. **a**. Schematic representation of a crRNA-Cas12a complex performing *trans-* cleavage of ssDNA reporters following the recognition of two split-activators. **b-d**. Fold change at t=60 minutes of *in vitro trans-*cleavage assay with Cas12a orthologs (red = LbCas12a, green = AsCas12a, orange = ErCas12a) activated by individual truncated ssDNA activators of length 6-20 nt **e-g**. Heat maps representing fold change at t=60 minutes of an *in vitro trans-*cleavage assay activated by combinations of truncated ssDNA activators of different lengths ranging from 6-14 nt in the Pp and Pd regions. The reactions contained 25 nM truncated ssDNA GFP-activators, 60 nM Cas12a, and 120 nM crGFP and were incubated for 60 min at 37°C. Error bars represent SD (n=3). Statistical analysis was performed using a two-tailed t-test where ns = not significant with p > 0.05, and the asterisks (* p ≤ 0.05, ** p ≤ 0.01,*** p ≤ 0.001, and **** p ≤ 0.0001) denote significant differences.

We first performed *in vitro trans-*cleavage assays with individual truncated activators using three different orthologs of Cas12a. We found that the *trans-*cleavage activity is extremely sensitive to truncations of the ssDNA activators across the tested orthologs (Fig. 1b-d). Compared to the full-length 20-nt activator, the *trans-*cleavage activity for a 16-nt activator diminishes by as much as 50-70-fold and is completely lost for activators less than 12-nt in length. Unlike LbCas12a and ErCas12a which show no *trans-*cleavage, AsCas12a shows a detectable activity for the target of length 14-nt. These results corroborate an earlier study that demonstrated a crRNA-target DNA interaction longer than 14-nt is necessary to initiate the indiscriminate *trans-*cleavage activity of Cas12a^34^.

Next, we tested to check if the simultaneous addition of two truncated activators, each binding to different regions of the crRNA and mimicking a full-length target, would be able to regain the lost *trans-*cleavage activity (Fig. S2). For this, we analyzed the activity produced by truncated activators shorter than 14-nt in a combinatorial fashion (Fig. 1e-g). We were surprised to observe that while the individual truncated activators failed to trigger any *trans-* cleavage activity, a split-activator combination was able to partially regain the diminished activity when the combined lengths were greater than or equal to 20-nt. This demonstrated that Cas12a enzymes can accept two different activators in a split-activator fashion and still initiate *trans*-cleavage.

### Cas12a tolerates RNA activators at the PAM-distal 3’-end of the crRNA

After observing the *trans-*cleavage behavior of Cas12a orthologs towards truncated ssDNA split-activators, we wondered how sensitive the Pd and Pp regions were to other nucleic acid substrates such as RNA and dsDNA. We were particularly curious to see how RNA substrates would be tolerated in the split-activator system since Cas12a is not known for initiating *trans-*cleavage after targeting RNA. To study this, we designed 10-nt ssDNA, dsDNA, and RNA activators complementary to either the Pp or the Pd region of the crRNA (Fig. 2a).

**Fig. 2.**
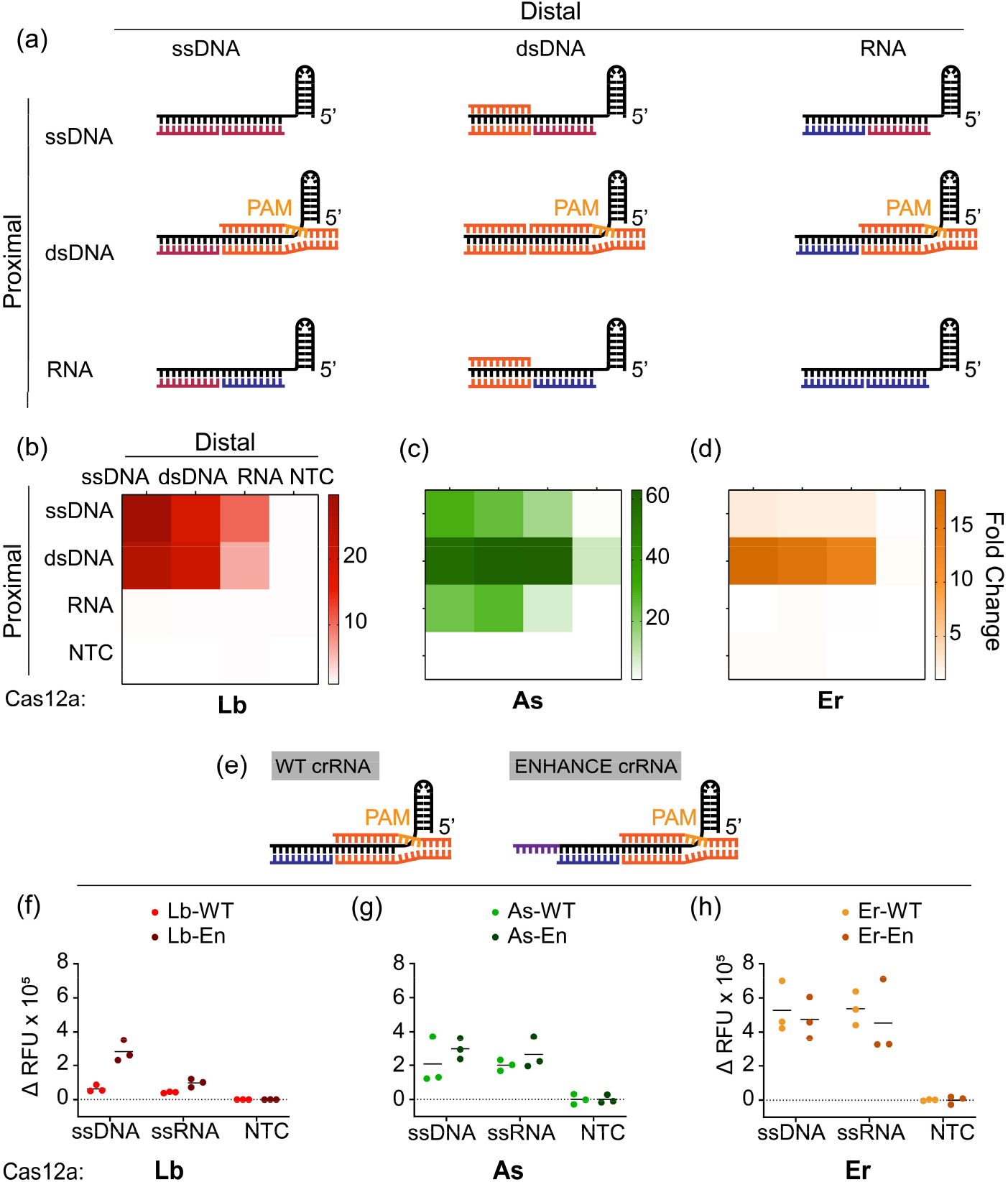
Split-activator detection of ssDNA, dsDNA, and RNA substrates by Cas12a: **a**. Schematic representation of Cas12a activated by combinations of ssDNA (red), dsDNA (orange), and RNA (blue) in the PAM proximal and PAM distal regions. **b-d** Heat maps representing the fold changes of *in vitro trans-*cleavage assay (n=3) with Cas12a orthologs for the combinatorial schemes seen in **(a). e-h** Comparison of the WT crRNA and ENHANCE crRNA for *in vitro trans-*cleavage assay split activators. Note, ssDNA and ssRNA substrates were used as targets in the Pd region while dsDNA was supplied in the Pp. Reactions were incubated for 60 min at 37°C. Error bars represent SD (n=3). The reactions contained 25 nM of each truncated activator, 60 nM Cas12a, and 120 nM crRNA.

We tested the detection of different ssDNA, dsDNA, and RNA activators in a combinatorial fashion (Fig. 2b-d). Upon switching from ssDNA to RNA at the Pp seed region, we observed that the *trans-*cleavage activity completely vanishes for Lb and Er irrespective of the type of substrate supplied at the Pd. This demonstrated a strict DNA substrate requirement at the seed region. On the contrary, AsCas12a seemed to tolerate even RNA substrates at the Pp region, hinting at distinct structural recognition features from other Cas12a orthologs. Interestingly, all three orthologs were observed to tolerate RNA substrates at the Pd region of the crRNA, provided a short piece of DNA substrate is supplied at the Pp region. The DNA binding at the Pp can be ssDNA or dsDNA. However, we observed that the *trans-*cleavage activity is significantly boosted for As and Er with a PAM-containing dsDNA at the Pp region instead of an ssDNA.

While PAM is not necessary for ssDNA target sequences, Cas12a is known to require a PAM sequence for targeting dsDNA. We speculated whether the PAM sequence is also critical for dsDNA binding in a split-activator manner. To test, we designed PAM-containing (+PAM) as well as PAM-deprived (-PAM) dsDNA activators complementary to either the Pp or the Pd (Fig. S3). We observed the split activator system to have the best activity when a +PAM dsDNA binds at the Pp and the −PAM dsDNA binds at the Pd. We also observed that there is no activity when a −PAM dsDNA binds at the Pp-end, indicating that a PAM is crucial for dsDNA binding at the Pp-end.

Previously, we have shown that adding a 7-nt DNA extension at the 3’-end of the crRNA significantly boosts the trans-cleavage activity of LbCas12a^22^. We were curious to check if these modified crRNAs (termed ENHANCE or EN) would have any significant effect on the *trans-*cleavage activity produced by the split-activator system. We analyzed the detection of ssDNA and ssRNA substrates using the split-activator system with both the wild-type (WT) and EN crRNAs. We observed that both ssDNA and ssRNA had significantly greater activity with the EN crRNA for LbCas12a, while AsCas12a and ErCas12a were unaffected by the different crRNA designs. This is consistent with previous observations with ENHANCE. We observed ErCas12a to have the highest activity with the split-activator system irrespective of the crRNA design being used. (Fig. 2f-h).

Chimeric DNA-RNA hybrid guides have been previously used to increase the sensitivity and reduce the off-target effects for both Cas9 and Cas12 nucleases^22,35,36^. We questioned if the use of a chimeric DNA-RNA hybrid crRNA would help with the detection of split-activator sequences. To investigate this, we designed two chimeric crRNAs by changing either 12-nt at the Pp region of the crRNA to DNA (crRNA-12D8R) or 8-nt of the Pd region of the crRNA to DNA (crRNA-12R8D) (Fig.S4). However, our results indicated that the chimeric DNA-RNA hybrid guides performed poorly in the detection of split activators (Fig. S5).

### Development of SAHARA for the detection of different lengths of RNA targets

Using our observations, we developed SAHARA (<u>S</u>plit <u>A</u>ctivators for <u>H</u>ighly <u>A</u>ccessible <u>R</u>NA <u>A</u>nalysis) a tool for direct RNA detection with Cas12a. The design of SAHARA was such that a short (12-nt), synthetic PAM-containing dsDNA activator (named ‘S12’ due to its length) would bind to the Pp-end of the crRNA, while any target RNA sequence can be detected at the Pd-end. While it was clear that short RNA activators could be easily detected by SAHARA, this study aimed to aid in the optimization of detection for a variety of targets and lengths. Therefore, we designed two RNA targets: a short RNA (20-nt) and a longer RNA (~730-nt) containing the same 20-nt target sequence (Fig. 3a).

**Fig. 3:**
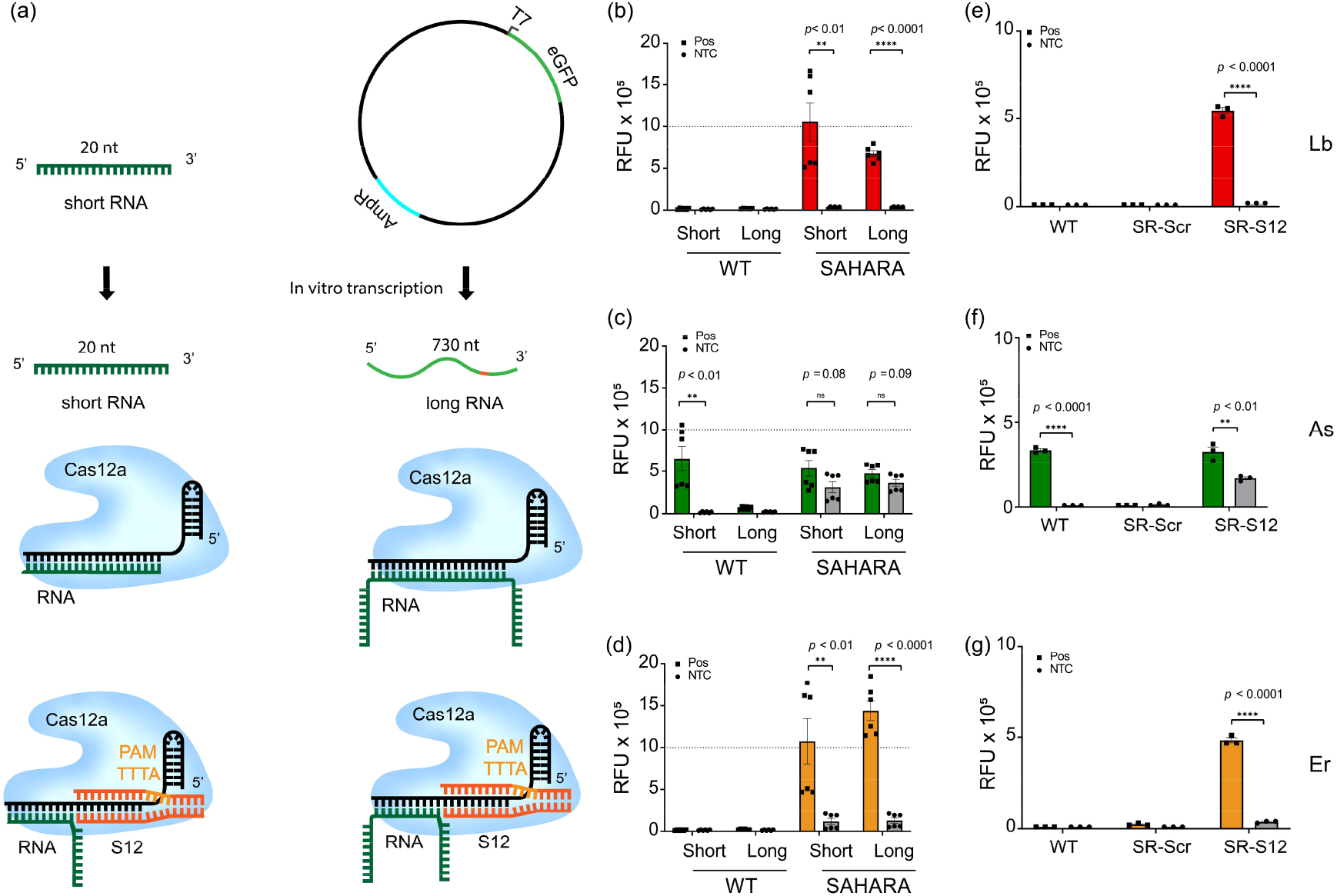
Development of SAHARA for the detection of a wide range of RNA targets. **a**. Schematic representation of Cas12a complexed with WT vs. SAHARA crRNA and activated by either a short (20-nt) or long (730-nt) RNA activators. **b-d**. Comparison of *trans-*cleavage activity among Cas12a orthologs for the short vs. long combinatorial schemes seen in **(a)**. The plot represents raw fluorescence units (RFU) plotted for time t=60 min. **e-g**. Detection of RNA target with SAHARA by using either a non-targeting scrambled S12 (SR-Scr) or a targeting S12 (SR-S12). The plot represents RFU at t=60 min. All error bars represent SD (n=3). Statistical analysis was performed using a two-tailed t-test where ns = not significant with p > 0.05, and the asterisks (* P ≤ 0.05, ** P ≤ 0.01,*** P ≤ 0.001, **** P ≤ 0.0001) denote significant differences.

To test these targets, we designed two crRNAs – a canonical WT as well as a SAHARA crRNA that only binds to the target at the Pd end while binding to a synthetic S12 at the Pp-end. We observed that while the WT CRISPR-Cas system failed, SAHARA was robustly able to detect RNA activators of different lengths (Fig. 3b-d). ErCas12a showed the best detection for both short and long RNA targets, while AsCas12a performed the worst. LbCas12a had decent activity for the short RNA target, but it performed significantly worse as the length of the RNA increased (Fig.3 a-d). We also observed that SAHARA does not show any RNA detection if a non-targeting ‘scrambled’ S12-activator is used instead of a canonical on-target S12, confirming S12-dependent activation of Cas12a activity (Fig.3 e-g).

Next, we studied whether the length of the crRNA has any effect on the RNA detection capability of SAHARA. We hypothesized that longer crRNAs would be better for RNA detection with SAHARA on account of their higher specificity of detection due to the increased number of base-pairing between the activator and the target. To test this, we designed crRNAs of length ranging from 24-28 nt. Each crRNA was complementary to their respective S12 activators at their Pp end while binding to an increasingly longer length of the target RNA at the Pd (Fig. S6). While all three crRNAs tested displayed some level of RNA detection, the 24-nt crRNA had the best activity for RNA detection with SAHARA.

While a fully complementary RNA activator is not recognized by Cas12a, we have shown that RNA can be recognized at the 3’-end of the crRNA if DNA is bound at the seed region. Therefore, somewhere between the seed region and the 3’-end of the crRNA protospacer, RNA recognition starts being tolerated. To examine the exact position at which RNA detection begins, we designed increasingly long RNA activators of length ranging from 6-14 nt that are complementary to the Pd region of the crRNA, and their corresponding DNA activators of different lengths binding to the Pp region (Fig. S7). We tested these DNA-RNA combinations in tandem and observed that while RNA activators of length 6-10 nt are recognized by Cas12a, increasing the length of the RNA activator beyond 10-nt stops any trans-cleavage activity. This suggests that the Pp 10-nt nucleotides of the crRNA protospacer can exclusively recognize DNA targets while the Pd 10-nt can detect both RNA and DNA.

While the catalytic activity of Cas12a enzymes is optimal at 37°C, the short (12-nt) DNA activators that are necessary for SAHARA to function, can bind more stably at temperatures lower than 37°C, due to their lower Tm. To study this, we tested the detection of an ssDNA and ssRNA target each at 37°C and room temperature (RT) (Fig. S8). We observed that both targets can be detected at RT. However, the activity drops by 3- to 5-fold depending on the Cas12a ortholog being used.

Cas12a effectors are metal-dependent endonucleases and therefore, the type and concentration of metal ions used in the reaction can have a significant effect on their activity. While Mg^2+^ ions are routinely used in Cas12a-based applications, Mn^2+^ ions have also been shown to work well^37^. To study the effect of different metal ions on the activity of SAHARA, we tested a range of different metal cations (NH_4_^+^, Rb^+^, Mg^2+^, Zn^2+^, Cu^2+^, Co^2+^, Ca^2+^, Ni^2+^, Mn^2+^, and Al^3+^) and discovered that most metal ions including severely inhibited the trans-cleavage activity of SAHARA (Fig.S9).

Among the different divalent metal ions tested, Mg^2+^ ions displayed the highest activity *in-vitro*, consistent with the literature. Therefore, we characterized the effect of increasing the concentration of Mg^2+^ ions on the activity. By varying the amount of Mg^2+^ ions in the Cas12a reaction, we observed a significant increase in activity with increasing concentrations up to 15 mM. Increasing the Mg^2+^ concentration beyond 15 mM slightly decreases the activity, suggesting 15 mM is the optimal concentration of Mg^2+^ for SAHARA (Fig. S10).

Finally, the pH of the reaction can have a notable effect on the charge of the enzyme and can subsequently affect its activity. We screened the activity of SAHARA in different buffers with pH ranging from 5.5-9.25. We observed that SAHARA has optimal activity at pH = 7.9 and the activity drops sharply upon further increasing or decreasing the pH (Fig. S11). There was no observed activity at pH = 9.25 for any of the three orthologs tested.

### Detection of HCV and miRNA-155 with SAHARA

To validate SAHARA with clinically relevant RNA targets, we designed multiple crRNAs targeting HCV RNA and miR-155. HCV is a viral infection that causes liver inflammation and can lead to serious complications such as liver damage, cirrhosis, and liver cancer. Early detection of HCV is important because it can help people receive treatment before the virus causes serious damage to the liver^38^. For HCV, we synthesized a short target RNA resembling a polypeptide precursor gene that is conserved across multiple HCV genotypes. For this RNA, we designed three different SAHARA guide RNAs each targeting it at either the 5’-end (head), 3’-end (tail), or in the middle (mid) (Fig. 4a).

**Fig. 4:**
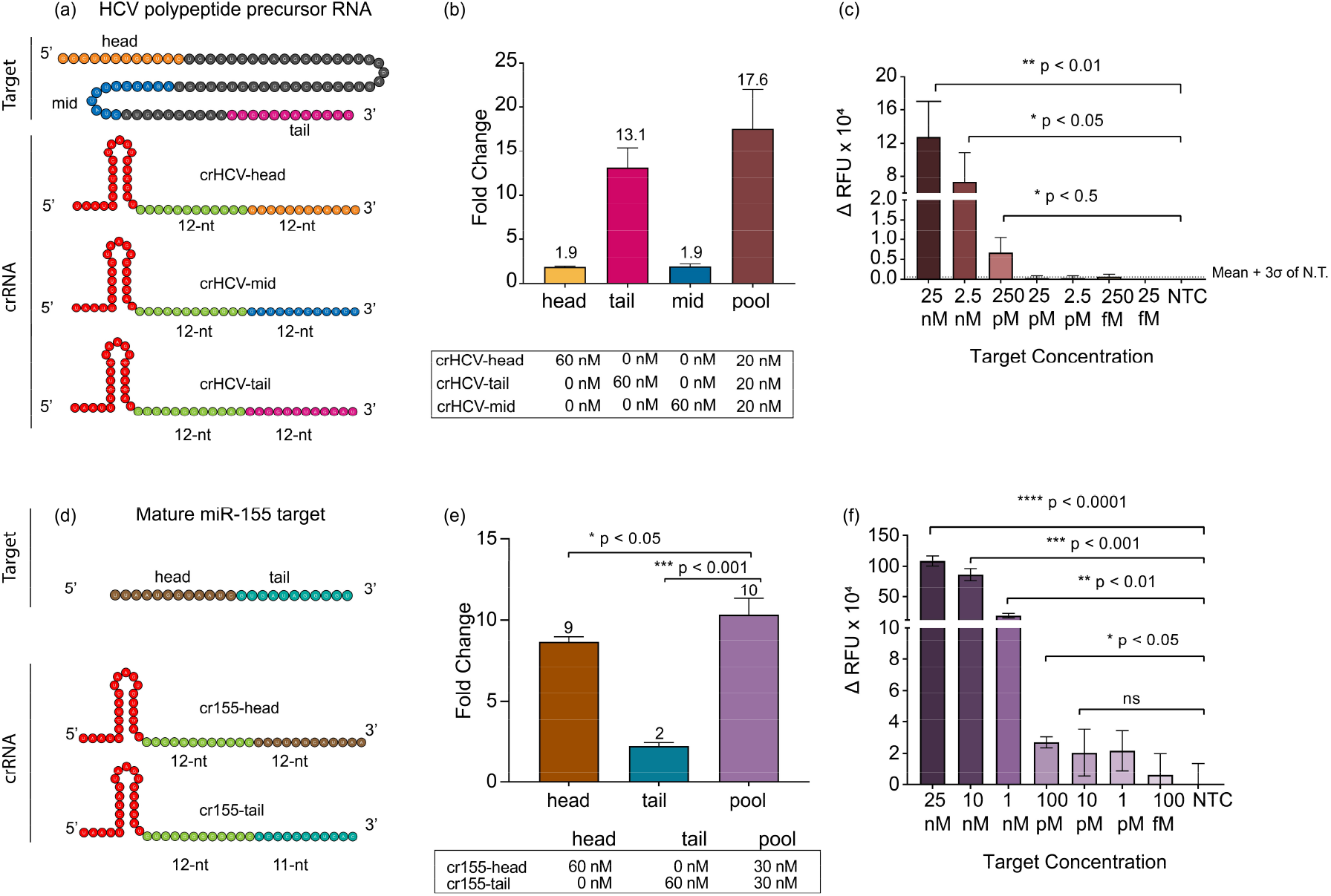
Application of SAHARA for the detection of HCV and miR-155. **a**. Schematic of an HCV polypeptide precursor RNA target and three crRNAs targeting it at different positions. Colors within the target and crRNA indicate the following: crRNA Pp region (green), head (orange), tail (purple), and middle (blue) sections of an HCV polypeptide precursor RNA. **b**. Comparison among the head, tail, mid, and pooled HCV targeting crRNA. The plot represents the fold change in fluorescence intensity normalized to the NTC at t=60 min (n=3). **c**. Limit of detection of HCV target using a pool of head, tail, and mid crRNA sequences. The plot represents the background-subtracted fluorescence intensity at t=60 min, for different concentrations of the target. **d**. Head vs Tail detection for a mature miRNA-155 target meditated by a split activator system. crRNAs were designed to target an S12 dsDNA GFP-activator in the Pp region and target either the head or tail region of a miR-155 target in the Pd region. **e**. Comparison of normalized fluorescence intensity fold change values among cr155-Tail, cr155-Head, and a combination of both Head and Tail targeting crRNAs. **f**. miR-155 limit of detection using a pooled crRNA with a split activator system. The plot represents the background-subtracted fluorescence intensity at t=60 min, for different concentrations of the target. Error bars represent SD (n=3). Statistical analysis was performed using a two-tailed t-test where ns = not significant with p > 0.05, and the asterisks (* P ≤ 0.05, ** P ≤ 0.01, *** P ≤ 0.001, **** P ≤ 0.0001) denote significant differences.

The miRNA-155 (or miR-155) is known to play a crucial role in breast cancer progression and is overexpressed in breast cancer tissues^39,40^. Therefore, reliable detection of miR-155 is important for early diagnosis. The mature miR-155 is ~23-nt in length. We synthesized the mature miRNA target and designed two SAHARA guides targeting it at either 12-nt at the 5’-end or 11-nt at the 3’-end, keeping the S12-binding region constant.

For both miR-155 and HCV targets, we first tested detection with the different designed guides individually. While all the designed guides for both targets were functional and displayed *trans-*cleavage, we observed significant variability in the activities produced by different guides. For miR-155, the head targeting guide showed an almost 5-fold higher activity than its tail targeting counterpart. Similarly, for HCV, the tail targeting guide showed as much as a 6-fold increase in activity as compared to head-targeting and the mid-targeting guide (Fig. 4b,e).

Upon closer inspection of the crRNAs with their corresponding targets, we observed that the secondary structure of the RNA target determines the activity of SAHARA (Fig. S12). Targets with a high amount of secondary structure are more inaccessible to bind to the Cas12a-crRNA complex, and are therefore harder to detect, while targets with relatively low or no secondary structure are detected easily. For miRNA-155, the fact that the tail guide only binds to 11-nt of the target as opposed to 12-nt binding in the head guide might play a role in the reduced activity.

It has previously been shown with Cas13a enzyme that pooling together multiple crRNAs enhances the level of detection^41^. We rationalized that a similar approach would also work with SAHARA. In concordance with the previous reports^41^, SAHARA also displayed a higher activity with pooled crRNAs for both miRNA-155 and HCV targets.

Finally, we tested to check the sensitivity of SAHARA. For this, we created dilutions of miRNA-155 and HCV targets ranging from 25 nM - 10 fM and tested for the detection of each target at different concentrations with their corresponding pooled crRNAs. We obtained a limit of detection of 132 pM for HCV and 767 pM for the miRNA-155 RNA target (Fig. 4c,f). These results are in concurrence with earlier studies showing similar limits of amplification-free detection of ssDNA and dsDNA with Cas12a^42^.

### SAHARA improves the specificity of target detection

We hypothesized that SAHARA might be more sensitive in discriminating single point mutations in the target as compared to the WT CRISPR-Cas12a since SAHARA binds to significantly fewer nucleotides of the target. To test this, we designed 12-nt ssDNA activators with single-point mutations for detection with SAHARA, as well as 24-nt ssDNA activators with identical mutations for detection of WT CRISPR-Cas12a for comparison (Fig. 5a,b). We observed that the detection of single-point mutants with both SAHARA and WT CRISPR was position-dependent compared to the WT target. We observed that mutations in positions M01-M06 significantly decreased the SAHARA-mediated *trans-*cleavage activity for Lb and Er (Fig. 5c-e). The same was true with positions M01-M04 with As but not with positions M05 and M06 (Fig. 5d). Mutations in positions beyond M06 did not decrease the *trans-*cleavage activity for either Cas12a ortholog. Our data indicated that SAHARA had improved specificity over WT-CRISPR for mutations near the split-activator boundary for the different Cas12a orthologs. This implied that mutations closer to the boundary of the split-activators have a larger effect on the *trans-*cleavage activity of SAHARA as compared to mutations away from the boundary.

**Fig. 5:**
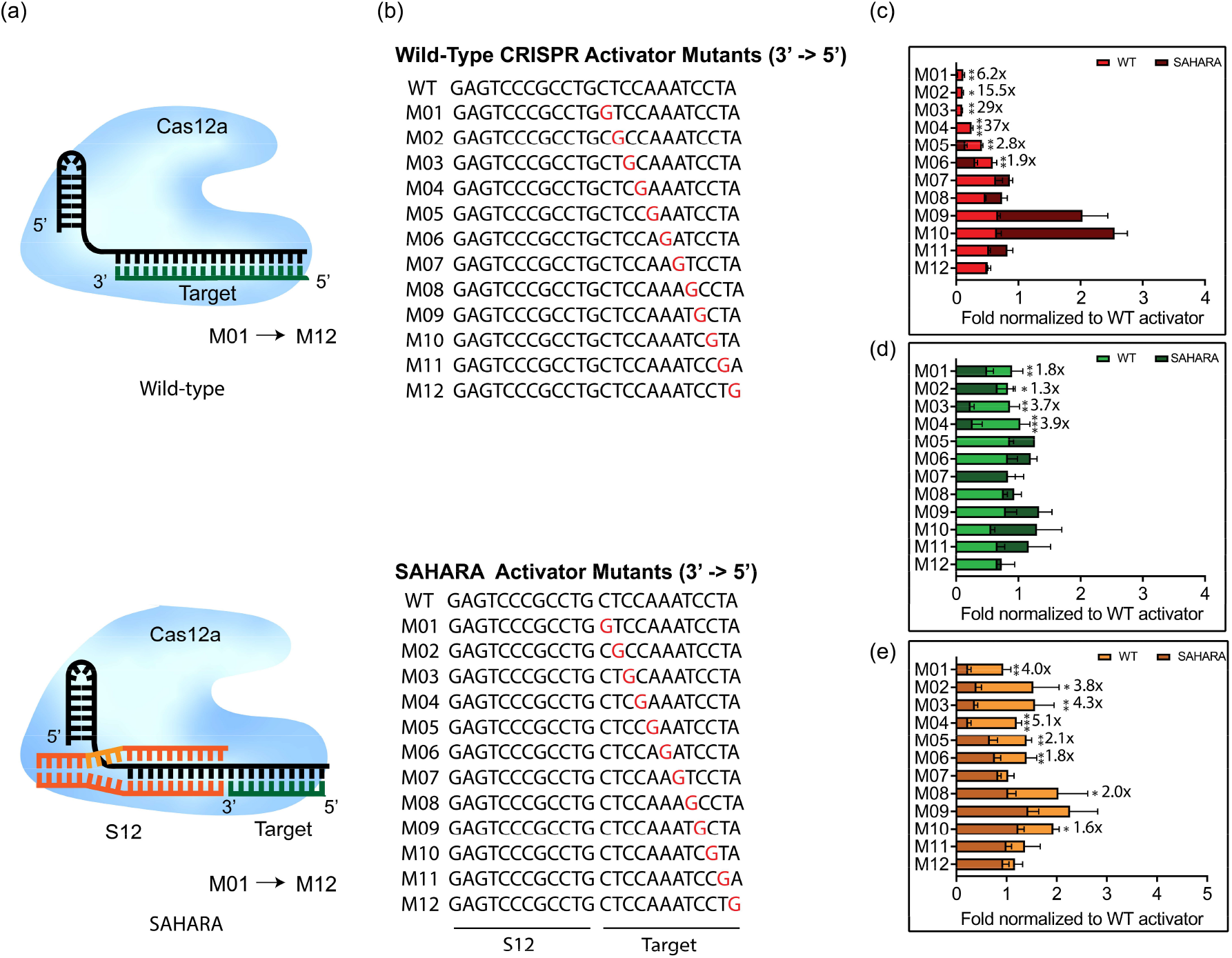
Specificity of SAHARA towards single point mutations in target. **a**. Schematic of WT vs SAHARA CRISPR-Cas systems for the detection of a target nucleic acid. **b**. ssDNA activators were designed with point mutations across the length of the activator. GFP-activator mutants were designed for a WT CRISPR activator (24-nt) and a SAHARA split activator system (12-nt +12-nt). The mutation location is identified by ‘M’ following the nucleotide number where the base has been changed to guanine (3’ to 5’ direction). **c-e**. Comparison of fold changes for the *in vitro trans-*cleavage assay between WT and SAHARA activator mutants normalized to the WT activator for Cas12a orthologs (c: LbCas12a, d: AsCas12a, and e: ErCas12a). Comparison of RFU values at t=60 min for the *in vitro trans-*cleavage assay between WT and SAHARA. Statistical analysis was performed using a two-tailed t-test where ns = not significant with p > 0.05, and the asterisks (* P ≤ 0.05, ** P ≤ 0.01,*** P ≤ 0.001, **** P ≤ 0.0001) denote significant differences.

### S12 DNA can be repurposed as a switch to control the *trans-*cleavage activity of SAHARA

To verify the PAM-dependency of SAHARA for initiating *trans-*cleavage, we designed three different S12 activators each containing either a TTTA, AAAT, or VVVN PAM respectively (Fig. 6a). TTTA is one of the canonical PAM for Cas12a, AAAT is the anti-PAM sequence, and VVVN encompasses the space of all the PAM sequences that are not tolerated by Cas12a. We tested the detection of a short 20-nt RNA with these S12 activators. As expected, the TTTA-PAM S12 was able to mediate RNA detection with all three Cas12a orthologs (Fig. 6b-d). Upon changing the PAM from TTTA to AAAT or VVVN, the *trans-*cleavage activity was completely diminished for Lb and As. Interestingly, Er Cas12a was able to detect RNA even with an AAAT or VVVN PAM, hinting at a broader PAM tolerance for *trans-*cleavage than previously reported.

**Fig. 6.**
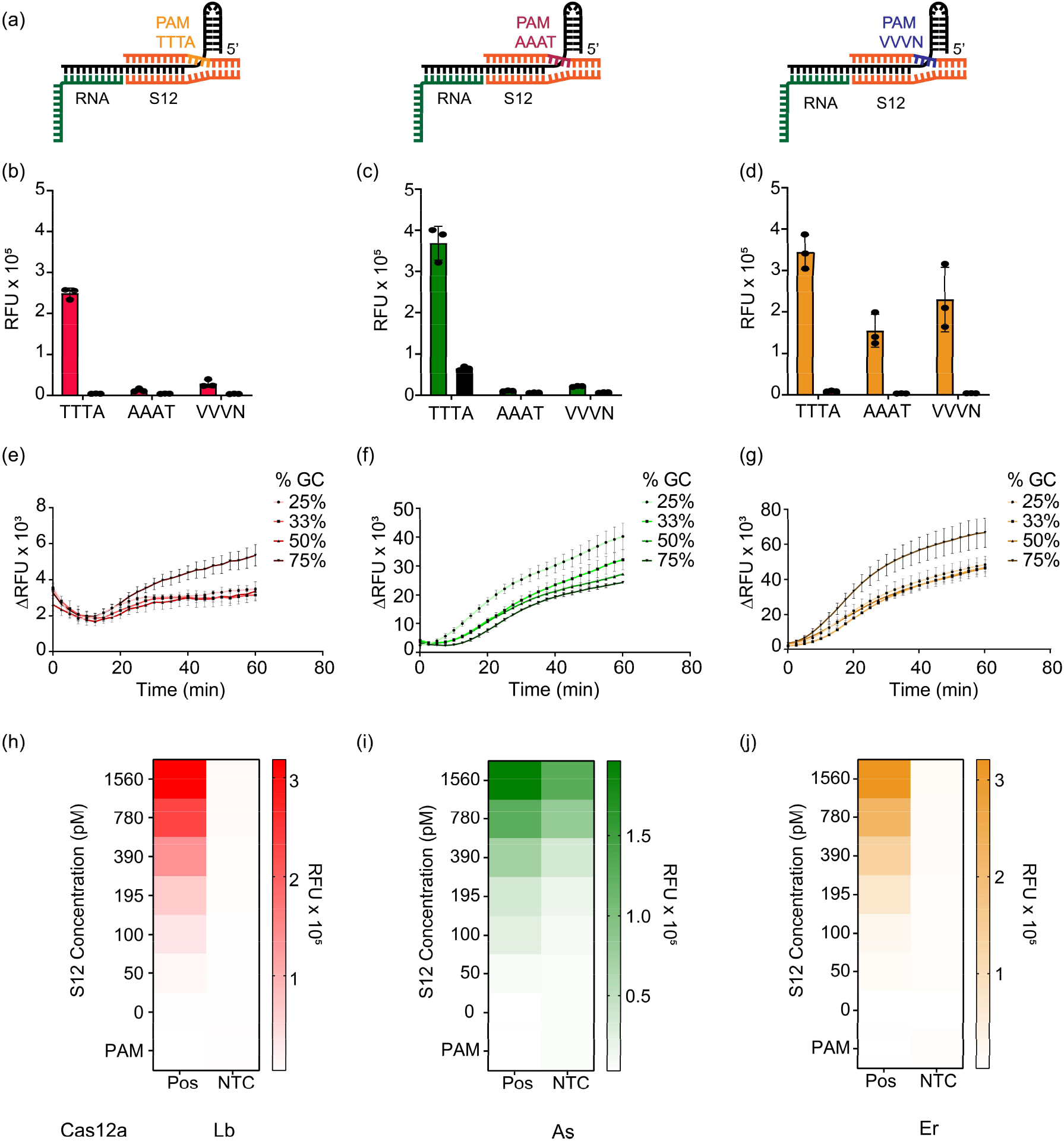
Characterizing properties of the S12 DNA binding at the Pp region in SAHARA: **a-d**. PAM sequence tolerance of Cas12a orthologs (red = LbCas12a, green = AsCas12a, orange = ErCas12a) coupled with SAHARA. Comparison of *trans-*cleavage activity among S12 dsDNA activators containing different PAM sequences (n=3). The PAM sequences TTTA, AAAT, and VVVN were assessed. **e-g**. Cas12a orthologs tolerate a wide range of GC contents in the crRNA and S12 dsDNA for RNA detection (n=3). **h-j**. The *trans-*cleavage activity of Cas12a with varying concentrations of S12 after incubation for 60 min at 37°C. Error bars represent SD (n=3).

**Fig. 7.**
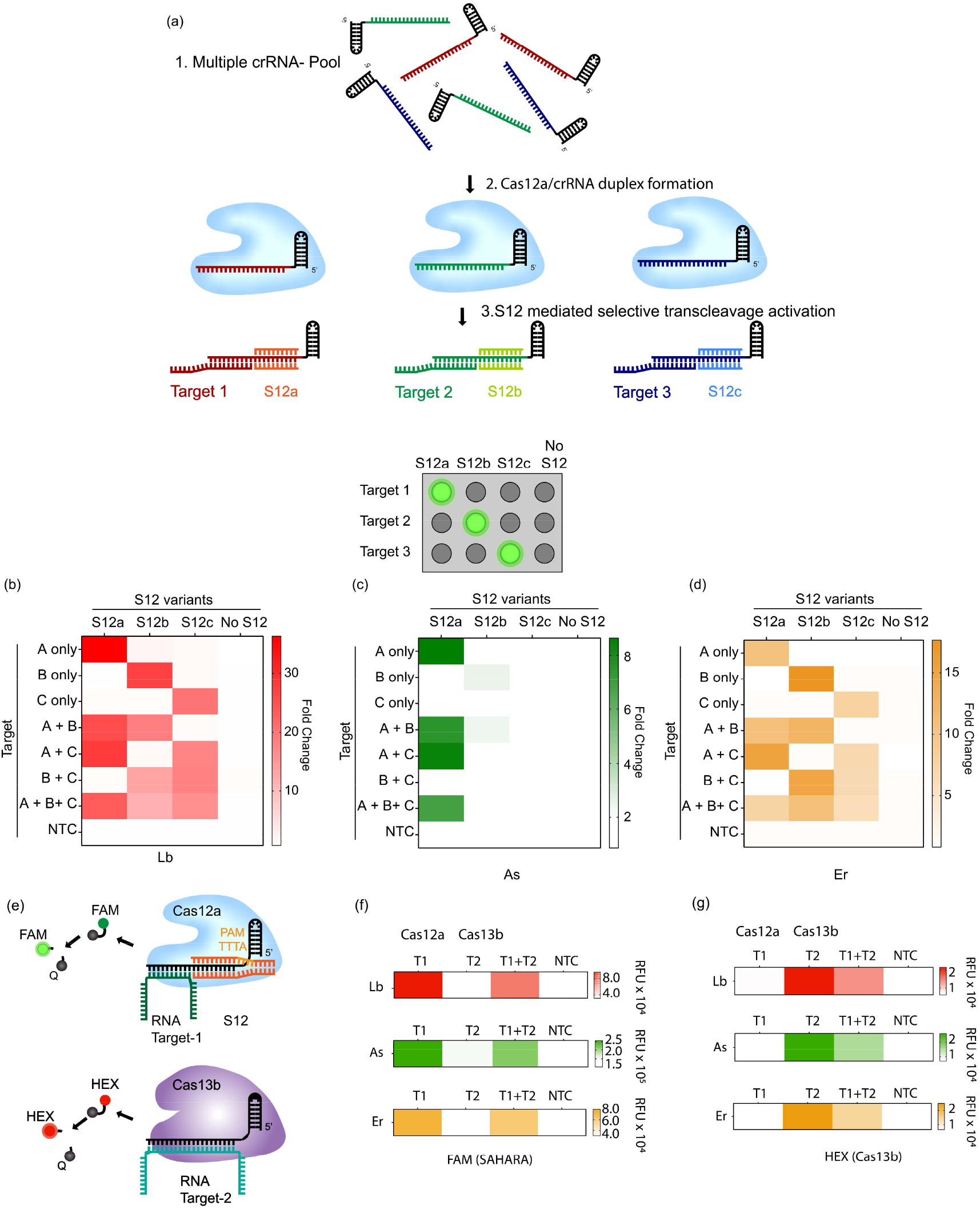
Simultaneous detection of multiple targets with SAHARA. **a**. Schematic of multiplexed detection with SAHARA. A mixture of different crRNAs can be differentiated for *trans-*cleavage activity by the use of sequence-specific S12 activators. **b-d**. Heat maps depicting the *trans-*cleavage activity of 3 different pooled crRNAs (crRNA-a, crRNA-b, and crRNA-c) in the presence of 3 different S12 activators (S12a, S12b, S12c) or a no S12 control for Lb, As, and Er cas12a orthologs. Fold change compared to the NTC at t=60 min from the start of the reaction is plotted. 30 nM Cas12a, 60 nM crRNA, 25 nM S12 activators, and 25 nM of DNA or RNA targets were used in the assay (n=3). **e**. Schematic of multiplexed RNA detection with a combination of SAHARA and Cas13b. DNA or RNA reporters consisting of different colored dyes are used to distinguish the signal produced by Cas12a and Cas13b. **f-g**. Multiplexed RNA detection using Lb, As, and Er orthologs of Cas12a and PsmCas13b. Cas12a targets activator T1 and produces a signal in the FAM channel, while Cas13b targets activator T2 and produces a signal in the HEX channel (n=3).

Next, we conjectured whether the GC content of the S12 activator plays a role in the RNA detection activity of SAHARA. To check this, we designed different crRNAs and S12 activators consisting of GC content ranging from 25-75%. All 3 Cas12a orthologs were able to tolerate changes in the GC content for the activity of SAHARA. This implies that Cas12a orthologs can tolerate a wide range of GC content for RNA detection.

Finally, we tested to check the minimum concentration of S12 needed to initiate RNA detection with SAHARA. We varied the concentration of S12 from 0-1.5 nM in increasing amounts and tested for the detection of a 25 nM RNA target. We observed an increase in the *trans-*cleavage activity of different Cas12a orthologs with increasing S12 concentration thereby suggesting that the activity of SAHARA is S12-dependent (Fig. 6h-j, S13, S14). Surprisingly, an S12 concentration as low as 50 pM concentration was sufficient to initiate *trans-*cleavage activity for a CRISPR-Cas12a complex consisting of 30 nM Cas12a and 60 nM crRNA. This suggests that a low amount of S12 is required to initiate the *trans-*cleavage activity of SAHARA. Remarkably, we observed that in the absence of S12, there was no *trans-*cleavage despite the presence of crRNA, Cas12a, and target. This suggested that S12 is critical for the initiation of *trans-*cleavage activity and can be used as a switch to selectively turn the activity ON or OFF.

### Multiplexed DNA and RNA detection with SAHARA

We could leverage the switch-like function of the seed-binding S12 DNA activator to selectively turn ON the *trans-*cleavage activity of an individual crRNA from a pool of multiple different crRNAs. To investigate this, we used three different crRNAs (crRNA-a, crRNA-b, and crRNA-c), with unique targets and S12 activators, and pooled them all together. To demonstrate that we can simultaneously detect DNA and RNA with SAHARA, we used an ssDNA target A for crRNA-a while targets B and C were ssRNAs. We then performed the detection of each of the three targets with the pooled crRNAs and different Cas12a orthologs, in a combinatorial fashion, in the presence or absence of the corresponding S12. We also used a control wherein no S12 sequence was supplied to the crRNA-Cas-target mix.

In the no S12 control, there was no *trans-*cleavage activity with any of the target combinations despite the crRNA-Cas complex and the target being mixed, further reinforcing the idea that S12 is critical for activity with SAHARA. In the presence of different S12s, only the mixtures with the corresponding target and crRNA displayed *trans-*cleavage. For instance, in the presence of S12a, only the reactions where target A was available (A only, A+B, and A+B+C) were active, whereas the reactions without target A (B only, C only, or B+C) were inactive despite the Cas, crRNA-b, crRNA-c, and their respective targets being mixed. The data here shows strong evidence that the S12 DNA can be used to selectively activate specific crRNAs and can be used to control the *trans-*cleavage activity of CRISPR-Cas12a, thereby enabling simultaneous and multiplexable detection of both DNA and RNA targets.

Next, we postulated if we could perform multiplexed RNA detection by combining SAHARA with Cas13b^26^. We used DNA and RNA reporters consisting of orthogonal dyes to differentiate the signal obtained from SAHARA and Cas13. This is similar to the multiplexed detection with Cas13 and Cas12 demonstrated before^26^, but here we aimed to detect RNA substrates with both Cas12 and Cas13, and not DNA. To test this, we used Lb, As, and Er Cas12a orthologs in conjunction with PsmCas13b to detect two distinct RNA targets (T1 and T2). We designed Cas12 and Cas13 guide RNAs such that the guide for Cas12 was complementary to only the T1 target while the guide for Cas13 was complementary to only T2. We also used a FAM-containing DNA reporter and a HEX-containing RNA reporter to distinguish the signals of SAHARA and Cas13b.

Upon performing the detection of the T1 and T2 targets individually and in conjunction, we observed that SAHARA produced *trans-*cleavage activity only in the presence of the T1 target, while Cas13b produced *trans-*cleavage only in the presence of T2 (Fig.7 f, g). Furthermore, the *trans-*cleavage signal obtained from SAHARA and Cas13b could be distinguished from each other by using orthogonal fluorescent dyes such as FAM and HEX on different types of reporter molecules. Thus, it is feasible to combine SAHARA with Cas13b for multiplexed detection of distinct RNA targets.

## Discussion

Engineering enzymes to improve their functionality above and beyond what they are naturally capable of has been a long-standing goal of molecular biology. Often, the engineering is done through man-made intervention after careful structural and biochemical analyses. Rarely, and even more excitingly, novel properties of already well-characterized enzymes are discovered, putting them under a new spotlight and giving them a fresh perspective. While CRISPR-Cas systems have been studied for almost a decade, *trans-*cleaving Cas enzymes such as Cas12 and Cas13 are relatively new.

Despite their novelty, tremendous progress is being made toward studying and understanding their underlying mechanism, especially due to their utility in molecular diagnostics. Cas12a has been extensively studied and is widely used in diagnostic platforms such as DETECTR and ENHANCE. However, Cas12a-based diagnostic methods are limited to DNA detection, since RNA substrates are not innately tolerated by Cas12a. Here we have discovered that substrate tolerance of Cas12a enzymes is position-dependent concerning the crRNA, and that RNA substrates can be used to turn on the *trans-*cleavage activity of Cas12a by binding at the PAM-distal end of the crRNA, as long as a DNA substrate is supplied at the PAM-proximal end.

In contrast to Cas9, which uses two different active sites to generate a blunt double-stranded DNA break^43^, Cas12a uses a single active site to make staggered cuts on the two strands of a dsDNA^6,12,16^. After cleaving the target DNA, Cas12a releases the PAM-distal cleavage product while retaining the PAM-proximal cleavage product bound to the crRNA^29,44^. This maintains Cas12a in a catalytically competent state, in which the active site of RuvC remains exposed to the solvent which then leads to *trans-*cleavage of neighboring single-stranded DNA molecules in a nonspecific manner. It has been shown that crRNA-target DNA hybrids of length 14-nt or less do not trigger any *cis*- or *trans*-cleavage activity, and a crRNA-DNA hybrid of at least 17-nt is crucial for stable Cas12a binding and cleavage. These observations suggest that the interaction of Cas12a with crRNA-target DNA hybrid at positions 14-17 nt is critical for initiating cleavage^29,34,45^.

We discovered that truncating the length of ssDNA activators binding to the crRNA drastically diminishes the *trans-*cleavage activity of Lb, As, and Er Cas12a. However, while individual truncated activators fail to initiate *trans-*cleavage, supplying two truncated activators, each binding to a different region of the crRNA in a split-activator fashion, partially regains the lost activity. We also observed that in the case of LbCas12a, the longer the length of the activator binding to the Pp region of the crRNA, the greater the recovery in the *trans-*cleavage activity. Thus, the activator combination of 14-nt (Pp) + 6-nt (Pd) shows higher activity than the combination of 6-nt (Pp) + 14-nt (Pd) despite having identical total length and sequence. These observations reinforce the idea that the seed region of the crRNA plays a vital role in modulating the *trans-*cleavage activity of Cas12a orthologs.

Furthermore, upon changing the truncated activators from ssDNA to dsDNA or RNA, we observed that while the PAM-proximal seed region of the crRNA exclusively tolerates DNA for initiating *trans-*cleavage, the PAM-distal end of the crRNA could tolerate RNA along with DNA. The exception here was AsCas12a, which was able to tolerate RNA even at the PAM-proximal end, unlike Lb or Er Cas12a. We also observed that using a PAM-containing dsDNA at the Pp-end greatly increases the *trans-*cleavage as compared to using an ssDNA for two of the three Cas12a orthologs (As and Er).

Surprised by our results, we wondered if we could detect longer lengths of RNA sequences at the PAM-distal end of the crRNA. To enable this, we designed long crRNAs containing 24-nt of the spacer. Within this spacer, 12-nt at the Pp end was complementary to a PAM-containing dsDNA that we call S12, while 12-nt at the distal end was complementary to the target of interest. We named this split-activator-based method of RNA detection with Cas12a as SAHARA. Upon testing the detection of short and long RNA substrates with WT CRISPR-Cas12a and SAHARA, we observed that WT Cas12a could not detect RNA at all, or in the case of As, had a very small level of RNA detection. With SAHARA, however, all three cas12a orthologs displayed robust RNA detection activity for different lengths of RNA. ErCas12a was observed to have the strongest activity for the detection of long RNA sequences.

Furthermore, we found that the presence of a PAM sequence on the S12 is necessary for robust *trans-*cleavage activity and that changing the PAM sequence eliminates the activity for Lb and As Cas12a while reducing it for Er. Changing the GC content of the S12 activators seemed to have negligible effects on the activity of all three orthologs, suggesting that a wide range of GC content can be used for the S12 activator. Remarkably, we found that the S12 concentration as low as 50 pM was sufficient to trigger the *trans-*cleavage of a 30 nM crRNA-Cas complex for detecting a 25 nM target. These results indicated that very low levels of the S12 DNA binding to the crRNA are sufficient for detection; however, the fluorescence intensity output is increased at higher concentrations. Notably, in the absence of the S12, there is no *trans-* cleavage activity with any of the three Cas12a orthologs, implying that the S12 can be used as a switch to control the *trans-*cleavage of Cas12a.

We applied SAHARA for the detection of clinically relevant targets such as Hepatitis C virus (HCV) and miRNA-155. We found that the secondary structure of the target played an important role in the RNA detection activity of Cas12a. Increased secondary structure in the RNA target makes it more inaccessible to bind to the crRNA and reduces the activity. We also observed that pooling together multiple crRNAs targeting different regions of the activator enhanced the detection capability, as previously reported for Cas13. Using this knowledge, we were able to detect picomolar levels of both miRNA-155 as well as HCV with SAHARA.

We also found SAHARA to be highly specific to mutations at certain positions along the target. Mutations at positions closer to the interface of the S12 DNA and the bound target seemed to be especially detrimental in inhibiting activity. Finally, we used the switch-like behavior of the S12 activator to turn ON the *trans-*cleavage activity of a single crRNA from a pool of different crRNAs. We were also able to combine SAHARA with Cas13b for multiplexed detection of different RNA targets using DNA and RNA reporters containing orthogonal fluorescent dyes. Thus, SAHARA can be used as a unique diagnostic system that can detect both DNA and RNA targets simultaneously and in a multiplexable fashion.

## Methods

### Plasmid construction

Plasmids expressing Lb, As, and ErCas12a enzymes were constructed as described in Nguyen et.al^46^. Briefly, plasmids expressing LbCas12a and AsCas12a were obtained from Addgene (a gift from Zhang lab and Doudna lab) and directly used for protein expression. For ErCas12a, a plasmid containing the human codon-optimized Cas12a gene was obtained from Addgene, then was PCR amplified using Q5 Hot Start high fidelity DNA polymerase (New England Biolabs, Catalog #M0493S), and subcloned into a bacterial expression vector (Addgene plasmid #29656, a gift from Scott Gradia). The product plasmids were then transformed into Rosetta™(DE3)pLysS Competent Cells (Millipore Sigma, Catalog #70956) following the manufacturer’s protocols.

### Protein expression and purification

For protein production, bacterial colonies containing the protein-expressing plasmid were plated on an agar plate and grown at 37°C overnight. Individual colonies were then picked and inoculated for 12 hours in 10 mL of LB Broth (Fisher Scientific, Catalog #BP9723-500). The culture was subsequently scaled up to a 1.5 L TB broth mix and grown until the culture reached an OD = 0.6 to 0.8. The culture was then placed on ice before the addition of Isopropyl β-d-1-thiogalactopyranoside (IPTG) to a final concentration of 0.5 mM. The culture was then continued to grow overnight at 16°C for 14-18 hours.

The overnight culture was pelleted by centrifuging at 10,000 x g for 5 minutes. The cells were then resuspended in lysis buffer (500 mM NaCl, 50 mM Tris-HCl, pH = 7.5, 20 mM Imidazole, 0.5 mM TCEP, 1 mM PMSF, 0.25 mg/mL Lysozyme, and DNase I). The cell mixture was then subjected to sonication followed by centrifugation at 39800 x g for 30 minutes. The cell lysate was filtered through a 0.22 µm syringe filter (Cytiva, Catalog #9913-2504) and then ran through into 5 mL Histrap FF (Cytiva, Catalog #17525501, Ni^2+^ was stripped off and recharged with Co^2+^) pre-equilibrated with Wash Buffer A (500 mM NaCl, 50 mM Tris-HCl, pH = 7.5, 20 mM imidazole, 0.5 mM TCEP) connected to BioLogic DuoFlow™ FPLC system (Bio-rad).

The column was eluted with Elution Buffer B (500 mM NaCl, 50 mM Tris-HCl, pH = 7.5, 250 mM imidazole, 0.5 mM TCEP). The eluted fractions were pooled together and transferred to a 10 kDa – 14 kDa MWCO dialysis bag. Homemade TEV protease (plasmid was obtained as a gift from David Waugh, Addgene #8827, and purified in-house)(44) was added to the bag, submerged in Dialysis Buffer (500 mM NaCl, 50 mM HEPES, pH 7, 5 mM MgCl_2_, 2 mM DTT) and dialyzed at 4°C overnight.

The protein mixture was taken out of the dialysis bag and concentrated down to around 10 mL using a 30 kDa MWCO Vivaspin® 20 concentrator. The concentrate was then equilibrated with 10 mL of Wash Buffer C (150 mM NaCl, 50 mM HEPES, pH = 7, 0.5 mM TCEP) before injecting into 1 mL Hitrap Heparin HP column pre-equilibrated with Wash Buffer C operated in the BioLogic DuoFlow™ FPLC system (Bio-rad). The protein was eluted from the column by running a gradient flow rate that exchanges Wash Buffer C and Elution Buffer D (2000 mM NaCl, 50 mM HEPES, pH = 7, 0.5 mM TCEP). Depending on how pure the protein samples were, additional size-exclusion chromatography may have been needed. In short, the eluted protein from the previous step was run through a HiLoad® 16/600 Superdex® (Cytiva, Catalog #28989335). Eluted fractions with the highest protein purity were selected, pooled together, concentrated using a 30 kDa MWCO Vivaspin® 20 concentrator, snap-frozen in liquid nitrogen, and stored at –80°C until use.

### Target DNA, RNA, and guide preparation

All DNA and RNA oligos as well as the chimeric DNA/RNA hybrid crRNAs were obtained from Integrated DNA Technologies (IDT). Single-stranded oligos were diluted in 1xTE buffer (10 mM Tris, 0.1 mM EDTA, pH 7.5). Complementary oligos for synthesizing dsDNA were first diluted in nuclease-free duplex buffer (30 mM HEPES, pH 7.5; 100 mM potassium acetate) and mixed in a 1:5 molar ratio of target:non-target strand. Both strands were then subjected to denaturation at 95°C for 4 mins and gradient cooling at a rate of 0.1 °C/s to 25°C. For generating the 730-nt long GFP target sequence, Addgene plasmid pCMV-T7-EGFP (BPK1098) (Addgene plasmid # 133962, a gift from Benjamin Kleinstiver) was obtained and PCR amplified using Q5 Hot Start high fidelity DNA polymerase (New England Biolabs, Catalog #M0493S) from position 376-1125. The PCR amplified product was *in-vitro* transcribed using the HiScribe T7 High Yield RNA synthesis kit (NEB #E2040S) following the manufacturer’s protocol. The transcribed product was treated with DNase I for 30 min at 37°C and then purified using RNA Clean and Concentrator Kit (Zymo Research #R1016).

### Preparation of metal ion buffers

The different metal ion buffers were prepared by first creating a master mix of the following components: 50 mM NaCl, 10 mM Tris-HCl, and 100 µg/ml BSA. To this master mix, chloride salts of different monovalent, divalent, and trivalent cations (NH_4_^+^, Rb^+^, Mg^2+^, Zn^2+^, Co^2+^, Cu^2+^, Ni^2+^, Ca^2+^, Mn^2+^, and Al^3+^) were diluted to a final concentration of 10 mM. The pH of the buffer was adjusted to 7.9 by adding 1M NaOH.

### CRISPR-Cas12a reaction for fluorescence-based detection

All fluorescence-based detection assays were carried out in a low-volume, flat-bottom, black 384 well-plate. The crRNA-Cas12a conjugates were assembled by mixing them in NEB 2.1 buffer and nuclease-free water followed by incubation at room temperature for 10 min. The assembled crRNA-Cas12a mixes were then added to 250-500 nM FQ reporter and the necessary concentration of the target activator in a 40-μL reaction volume. The 384 well-plate was then incubated in a BioTek Synergy fluorescence plate reader at 37°C for 1 hour. Fluorescence intensity measurements for a FAM reporter were measured at the excitation/emission wavelengths of 483/20 nm and 530/20 nm every 2.5 min. A final concentration of 30 nM Cas12a, 60 nM crRNA, and 25 nM of target activator are used in all the assays unless otherwise specified.

### Limit of detection calculation

To find the limit of detection (LoD), the *trans-*cleavage assay was carried out with several different dilutions of the activator. The LoD calculations were based on the following formula^47^:

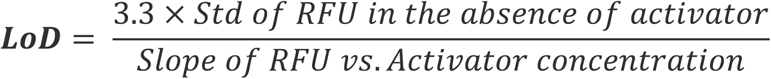

## Data Availability

All the data supporting the findings of this study are available within the Article and Supplementary Files or can be obtained from the corresponding author, P.K.J., upon reasonable request. Source data are available in the Source Data file. Source data are provided in this paper.

## Author Contributions

P.K.J., S.R.R., and S.S.A. conceived the idea and designed the experiments. S.R.R., E.K.V., G.M.S., S.S.A., L.S.S., K.S.M., and L.T.N. performed the experiments and data analysis. S.R.R., E.K.V., and G.M.S. wrote the initial manuscript. The manuscript was edited by P.K.J. and revised by all members.

## Acknowledgments

We are grateful to the members of the Jain lab for their helpful discussions and to the University of Florida (UF) Health Cancer Center for their support. Special thanks to Dr. Gary Wang and his lab members at the University of Florida for their insightful feedback on the project and Ms. Lilia G. Yang for her help with experiments and proofreading the manuscript. This work was financially supported by funds from the UF, the UF Herbert Wertheim College of Engineering, Shah Foundation Endowment Funds, Florida Breast Cancer Foundation (AGR00018466), NIH-NIAID R21AI156321, NIH-NIAID R21AI168795, and NIH-NIGMS R35GM147788. The funding sources did not have a role in the design of the study, the collection, analysis, or interpretation of data, nor in writing the manuscript.

