## Supplementary Information for "Programmable RNA detection with CRISPR-Cas12a"

Table S1: List of crRNA used in this study (5'→3'). The spacer region of SAHARA crRNAs is colored to indicate the positions bound by S12 activators (blue) and the target DNA or RNA (purple):

| Name | Sequence | Figure |
|---|---|---|
| crGFP-3'DNA7 (ENHANCE) | UAAUUUCUACUAAGUGUAGAUCUC AGGGCGGACUGGGUGCUTATTATT | Fig. 1b-g, Fig. 2b-h |
| crGFP-WT | UAAUUUCUACUAAGUGUAGAUCUC AGGGCGGACUGGGUGCU | Fig. 2f-h, Fig. 3b-h, |
| crGFP-SH | UAAUUUCUACUAAGUGUAGAUGA UUAGCAUUAACUCAGGGCGGAC | Fig. 3b-g, Fig. 7b-d |
| cr155-Head-SH | UAAUUUCUACUAAGUGUAGAUCUC AGGGCGGACGAUUAGCAUUAA | Fig. 4e,f |
| cr155-Tail-SH | UAAUUUCUACUAAGUGUAGAUCUC AGGGCGGACACCCCUAUCAC | Fig. 4e,f |
| cr155-Head-SH v2 | UAAUUUCUACUAAGUGUAGAUAU GGUGAGCAAGGAUUAGCAUUAA | Fig. 7b-d |
| crHCV-Head-SH | UAAUUUCUACUAAGUGUAGAUCUC AGGGCGGACGUACCACAAGGC | Fig. 4b,c |
| crHCV-Mid-SH | UAAUUUCUACUAAGUGUAGAUCUC AGGGCGGACGAUGCACGGUCU | Fig. 4b,c |
| crHCV-Tail-SH | UAAUUUCUACUAAGUGUAGAUCUC AGGGCGGACGAGGUUUAGGAU | Fig. 4b,c, Fig. 5c-e, Fig. 6e-g, Fig. 6h-j, Fig. 7b-d, Fig. 7f,g |
| crHCV-Tail-SH-25%GC | UAAUUUCUACUAAGUGUAGAUAUC AUAUCUGAAGAGGUUUAGGAU | Fig. 6e-g |
| crHCV-Tail-SH-33%GC | UAAUUUCUACUAAGUGUAGAUAUC AGAUCUGAAGAGGUUUAGGAU | Fig. 6e-g |
| crHCV-Tail-SH-50%GC | UAAUUUCUACUAAGUGUAGAUCUC AGAUCGGAAGAGGUUUAGGAU | Fig. 6e-g |

Table S2: List of Target Activators used in this study (5'→3'). The mutated positions for single-point mutants of HCV are indicated in red:

| Name | Sequence | Figure |
|---|---|---|
| GFP-20-nt | AGCACCCAGTCCGCCCTGAG | Fig. 1b-d |
| GFP-20-nt RNA | AGCACCCAGUCCGCCCUGAG | Fig. 3b-g, Fig. 7b-d |
| GFP-Pp-6-nt | CCTGAG | Fig. 1b-g |
| GFP-Pp-8-nt | GCCCTGAG | Fig. 1b-g |
| GFP-Pp-10-nt | CCGCCCTGAG | Fig. 1b-g, Fig. 2b-d |
| GFP-Pp-12-nt | GTCCGCCCTGAG | Fig. 1b-g |
| GFP-Pp-14-nt | CAGTCCGCCCTGAG | Fig. 1b-g |
| GFP-Pp-16-nt | CCCAGTCCGCCCTGAG | Fig. 1b-d |
| GFP-Pp-18-nt | CACCCAGTCCGCCCTGAG | Fig. 1b-d |
| GFP-Pd-6-nt | AGCACC | Fig. 1b-g |
| GFP-Pd-8-nt | AGCACCCA | Fig. 1b-g |
| GFP-Pd-10-nt | AGCACCCAGT | Fig. 1b-g, Fig. 2b-d, Fig. 2f-h |
| GFP-Pd-12-nt | AGCACCCAGTCC | Fig. 1b-g |
| GFP-Pd-14-nt | AGCACCCAGTCCGC | Fig. 1b-g |
| GFP-Pd-16-nt | AGCACCCAGTCCGCCC | Fig. 1b-d |
| GFP-Pd-18-nt | AGCACCCAGTCCGCCCTG | Fig. 1b-d |
| GFP-Pp-dsDNA-10-nt | CCGCCCTGAGTAAAGCG (TS)<br>GCGAAATGAGTCCCGCC (NTS) | Fig. 2b-d, Fig. 2f-h |
| GFP-Pd-dsDNA-10-nt | AGCACCCAGTTAAAGCG (TS)<br>GCGAAATTGACCCACGA (NTS) | Fig. 2b-d |
| GFP-Pp-RNA-10-nt | CCGCCCUGAG | Fig. 2b-d, |
| GFP-Pd-RNA-10-nt | AGCACCCAGU | Fig. 2b-d, Fig. 2f-h |

| | | |
|---|---|---|
| HCV polypeptide precursor RNA | GCCUUGUGGUACUGCCUGAUAGGGUGCUU GCGAGUGCCCCGGGAGGUCUCGUAGACCG UGCAUCAUGAGCACAAAUCCUAAACCUC | Fig. 4b,c, Fig.6 b-j, Fig. 7b-d, Fig. 7f,g |
| miRNA-155 target RNA | UUAAUGCUAAUCGUGAUAGGGGU | Fig. 4e,f, Fig. 7 b-d |
| GFP-RNA 730-nt | GAGAGCCGCCACCAUGGUGAGCAAGGGCG AGGAGCUGUUCACCGGGGUGGUGCCCAUC CUGGUCGAGCUGGACGGCGACGUAAACGG CCACAAGUUCAGCGUGUCCGGCGAGGGCG AGGGCGAUGCCACCUACGGCAAGCUGACC CUGAAGUUCAUCUGCACCACCGGCAAGCU GCCCGUGCCCUGGCCCACCCUCGUGACCA CCCUGACCUACGGCGUGCAGUGCUUCAGC CGCUACCCCGACCACAUGAAGCAGCACGA CUUCUUCAAGUCCGCCAUGCCCGAAGGCU ACGUCCAGGAGCGCACCAUCUUCUUCAAG GACGACGGCAACUACAAGACCCGCGCCGA GGUGAAGUUCGAGGGCGACACCCUGGUGA ACCGCAUCGAGCUGAAGGGCAUCGACUUC AAGGAGGACGGCAACAUCCUGGGGCACAA GCUGGAGUACAACUACAACAGCCACAACG UCUAUAUCAUGGCCGACAAGCAGAAGAAC GGCAUCAAGGUGAACUUCAAGAUCCGCCA CAACAUCGAGGACGGCAGCGUGCAGCUCG CCGACCACUACCAGCAGAACACCCCCAUC GGCGACGGCCCCGUGCUGCUGCCCGACAA CCACUACCUGAGCACCCAGUCCGCCCUGA GCAAAGACCCCAACGAGAAGCGCGAUCAC AUGGUCCUGCUGGAGUUCGUGACCGCCGC CGGGAUCACUCUCGGCAUGGACGAGCUGU ACAAG | Fig. 3b-d |
| HCV full-length WT | GAGTCCCGCCTGCTCCAAATCCTA | Fig. 5c-e |
| HCV full-length M01 | GAGTCCCGCCTG**G**TCCAAATCCTA | Fig. 5c-e |
| HCV full-length M02 | GAGTCCCGCCTGC**G**CCAAATCCTA | Fig. 5c-e |
| HCV full-length M03 | GAGTCCCGCCTGCT**G**CAAATCCTA | Fig. 5c-e |
| HCV full-length M04 | GAGTCCCGCCTGCTC**G**AAATCCTA | Fig. 5c-e |
| HCV full-length M05 | GAGTCCCGCCTGCTCC**G**AATCCTA | Fig. 5c-e |

| | | |
|---|---|---|
| HCV full-length M06 | GAGTCCCGCCTGCTCCA**G**ATCCTA | Fig. 5c-e |
| HCV full-length M07 | GAGTCCCGCCTGCTCCAA**G**TCCTA | Fig. 5c-e |
| HCV full-length M08 | GAGTCCCGCCTGCTCCAAA**G**CCTA | Fig. 5c-e |
| HCV full-length M09 | GAGTCCCGCCTGCTCCAAAT**G**CTA | Fig. 5c-e |
| HCV full-length M10 | GAGTCCCGCCTGCTCCAAATC**G**TA | Fig. 5c-e |
| HCV full-length M11 | GAGTCCCGCCTGCTCCAAATCC**G**A | Fig. 5c-e |
| HCV full-length M12 | GAGTCCCGCCTGCTCCAAATCCT**G** | Fig. 5c-e |
| HCV SAHARA WT | CTCCAAATCCTA | Fig. 5c-e |
| HCV SAHARA M01 | **G**TCCAAATCCTA | Fig. 5c-e |
| HCV SAHARA M02 | C**G**CCAAATCCTA | Fig. 5c-e |
| HCV SAHARA M03 | CT**G**CAAATCCTA | Fig. 5c-e |
| HCV SAHARA M04 | CTC**G**AAATCCTA | Fig. 5c-e |
| HCV SAHARA M05 | CTCC**G**AATCCTA | Fig. 5c-e |
| HCV SAHARA M06 | CTCCA**G**ATCCTA | Fig. 5c-e |
| HCV SAHARA M07 | CTCCAA**G**TCCTA | Fig. 5c-e |
| HCV SAHARA M08 | CTCCAAA**G**CCTA | Fig. 5c-e |
| HCV SAHARA M09 | CTCCAAAT**G**CTA | Fig. 5c-e |
| HCV SAHARA M10 | CTCCAAATC**G**TA | Fig. 5c-e |

| | | |
|---|---|---|
| HCV SAHARA M11 | CTCCAAATCC**G**A | Fig. 5c-e |
| HCV SAHARA M12 | CTCCAAATCCT**G** | Fig. 5c-e |

Table S3: List of 'seed-region' binding S12-activators used in this study (5'→3'):

| Name | Sequence | Figure |
|---|---|---|
| S12 for crGFP-SH | TTAATGCTAATCTAAAGCG (TS)<br>CGCTTTAGATTAGCATTAA (NTS) | Fig. 3b-g, Fig. 7b-d |
| S12 for crHCV-Head-SH | GTCCGCCCTGAGTAAAGCGA (TS)<br>TCGCTTTACTCAGGGCGGAC (NTS) | Fig. 4b,c |
| S12 for crHCV-Tail-SH | GTCCGCCCTGAGTAAAGCGA (TS)<br>TCGCTTTACTCAGGGCGGAC (NTS) | Fig. 4b,c, Fig. 6b-j, Fig. 6h-j, Fig. 7b-d, Fig. 7f,g |
| S12 for crHCV-Mid-SH | GTCCGCCCTGAGTAAAGCGA (TS)<br>TCGCTTTACTCAGGGCGGAC (NTS) | Fig. 4b,c |
| S12 for cr155-Head-SH | GTCCGCCCTGAGTAAAGCGA (TS)<br>TCGCTTTACTCAGGGCGGAC (NTS) | Fig. 4e,f |
| S12 for cr155-Tail-SH | GTCCGCCCTGAGTAAAGCGA (TS)<br>TCGCTTTACTCAGGGCGGAC (NTS) | Fig. 4e,f |
| S12 for cr155-Head v2 SH | CTTGCTCACCATTAAACAC (TS)<br>GTGTTTAATGGTGAGCAAG (NTS) | Fig. 6b-d |
| S12 for crHCV-Tail 25% | TTCAGATATGATTAAACAC (TS)<br>GTGTTTAATCATATCTGAA (NTS) | Fig. 6e-g |
| S12 for crHCV-Tail 33% | TTCAGATCTGATTAAACAC (TS)<br>GTGTTTAATCAGATCTGAA (NTS) | Fig. 6e-g |
| S12 for crHCV-Tail 50% | TTCCGATCTGAGTAAACAC (TS)<br>GTGTTTACTCAGATCGGAA (NTS) | Fig. 6e-g |
| S12 AAAT PAM | TTAATGCTAACATTTGCG (TS)<br>CGCAAATGTTAGCATTAA (NTS) | Fig. 6b-d |
| S12 VVVN PAM | TTAATGCTAACNBBBGCG (TS)<br>CGCVVVNGTTAGCATTAA (NTS) | Fig. 6b-d |

Table S4: List of protein sequences used in this study:

| **Name** | **Sequence** |
|---|---|
| LbCas12a | MSKLEKFTNCYSLSKTLRFKAIPVGKTQENIDNKRLLVEDEKRAEDYKGVK<br>KLLDRYYLSFINDVLHSIKLKNLNNYISLFRKKTRTEKENKELENLEINLRKEI<br>AKAFKGNEGYKSLFKKDIIETILPEFLDDKDEIALVNSFNGFTTAFTGFFDNRE<br>NMFSEEAKSTSIAFRCINENLTRYISNMDIFEKVDAIFDKHEVQEIKEKILNSD<br>YDVEDFFEGEFFNFVLTQEGIDVYNAIIGGFVTESGEKIKGLNEYINLYNQKT<br>KQKLPKFKPLYKQVLSDRESLSFYGEGYTSDEEVLEVFRNTLNKNSEIFSSIK<br>KLEKLFKNFDEYSSAGIFVKNGPAISTISKDIFGEWNVIRDKWNAEYDDIHLK<br>KKAVVTEKYEDDRRKSFKKIGSFSLEQLQEYADADLSVVEKLKEIIIQKVDEI<br>YKVYGSSEKLFDADFVLEKSLKKNDAVVAIMKDLLDSVKSFENYIKAFFGE<br>GKETNRDESFYGDFVLAYDILLKVDHIYDAIRNYVTQKPYSKDKFKLYFQNP<br>QFMGGWDKDKETDYRATILRYGSKYYLAIMDKKYAKCLQKIDKDDVNGN<br>YEKINYKLLPGPNKMLPKVFFSKKWMAYYNPSEDIQKIYKNGTFKKGDMFN<br>LNDCHKLIDFFKDSISRYPKWSNAYDFNFSETEKYKDIAGFYREVEEQGYKV<br>SFESASKKEVDKLVEEGKLYMFQIYNKDFSDKSHGTPNLHTMYFKLLFDEN<br>NHGQIRLSGGAELFMRRASLKKEELVVHPANSPIANKNPDNPKKTTTLSYDV<br>YKDKRFSEDQYELHIPIAINKCPKNIFKINTEVRVLLKHDDNPYVIGIDRGERN<br>LLYIVVVDGKGNIVEQYSLNEIINNFNGIRIKTDYHSLLDKKEKERFEARQNW<br>TSIENIKELKAGYISQVVHKICELVEKYDAVIALEDLNSGFKNSRVKVEKQVY<br>QKFEKMLIDKLNYMVDKKSNPCATGGALKGYQITNKFESFKSMSTQNGFIF<br>YIPAWLTSKIDPSTGFVNLLKTKYTSIADSKKFISSFDRIMYVPEEDLFEFALD<br>YKNFSRTDADYIKKWKLYSYGNRIRIFRNPKKNNVFDWEEVCLTSAYKELF<br>NKYGINYQQGDIRALLCEQSDKAFYSSFMALMSLMLQMRNSITGRTDVDFLI<br>SPVKNSDGIFYDSRNYEAQENAILPKNADANGAYNIARKVLWAIGQFKKAE<br>DEKLDKVKIAISNKEWLEYAQTSVKH |
| AsCas12a | MTQFEGFTNLYQVSKTLRFELIPQGKTLKHIQEQGFIEEDKARNDHYKELKPI<br>IDRIYKTYADQCLQLVQLDWENLSAAIDSYRKEKTEETRNALIEEQATYRNA<br>IHDYFIGRTDNLTDAINKRHAEIYKGLFKAELFNGKVLKQLGTVTTTEHENA<br>LLRSFDKFTTYFSGFYENRKNVFSAEDISTAIPHRIVQDNFPKFKENCHIFTRLI<br>TAVPSLREHFENVKKAIGIFVSTSIEEVFSFPFYNQLLTQTQIDLYNQLLGGISR<br>EAGTEKIKGLNEVLNLAIQKNDETAHIIASLPHRFIPLFKQILSDRNTLSFILEEF<br>KSDEEVIQSFCKYKTLLRNENVLETAEALFNELNSIDLTHIFISHKKLETISSAL<br>CDHWDTLRNALYERRISELTGKITKSAKEKVQRSLKHEDINLQEIISAAGKEL<br>SEAFKQKTSEILSHAHAALDQPLPTTLKKQEEKEILKSQLDSLLGLYHLLDWF<br>AVDESNEVDPEFSARLTGIKLEMEPSLSFYNKARNYATKKPYSVEKFKLNFQ<br>MPTLASGWDVNKEKNNGAILFVKNGLYYLGIMPKQKGRYKALSFEPTEKTS<br>EGFDKMYYDYFPDAAKMIPKCSTQLKAVTAHFQTHTTPILLSNNFIEPLEITK<br>EIYDLNNPEKEPKKFQTAYAKKTGDQKGYREALCKWIDFTRDFLSKYTKTTS<br>IDLSSLRPSSQYKDLGEYYAELNPLLYHISFQRIAEKEIMDAVETGKLYLFQIY |

| | |
|---|---|
| | NKDFAKGHHGKPNLHTLYWTGLFSPENLAKTSIKLNGQAELFYRPKSRMKR<br>MAHRLGEKMLNKKLKDQKTPIPDTLYQELYDYVNHRLSHDLSDEARALLPN<br>VITKEVSHEIIKDRRFTSDKFFFHVPITLNYQAANSPSKFNQRVNAYLKEHPET<br>PIIGIDRGERNLIYITVIDSTGKILEQRSLNTIQQFDYQKKLDNREKERVAARQ<br>AWSVVGTIKDLKQGYLSQVIHEIVDLMIHYQAVVVLENLNFGFKSKRTGIAE<br>KAVYQQFEKMLIDKLNCLVLKDYPAEKVGGVLNPYQLTDQFTSFAKMGTQ<br>SGFLFYVPAPYTSKIDPLTGFVDPFVWKTIKNHESRKHFLEGFDFLHYDVKTG<br>DFILHFKMNRNLSFQRGLPGFMPAWDIVFEKNETQFDAKGTPFIAGKRIVPVI<br>ENHRFTGRYRDLYPANELIALLEEKGIVFRDGSNILPKLLENDDSHAIDTMVA<br>LIRSVLQMRNSNAATGEDYINSPVRDLNGVCFDSRFQNPEWPMDADANGAY<br>HIALKGQLLLNHLKESKDLKLQNGISNQDWLAYIQELRN |
| ErCas12a | NNGTNNFQNFIGISSLQKTLRNALIPTETTQQFIVKNGIIKEDELRGENRQILK<br>DIMDDYYRGFISETLSSIDDIDWTSLFEKMEIQLKNGDNKDTLIKEQTEYRKA<br>IHKKFANDDRFKNMFSAKLISDILPEFVIHNNNYSASEKEEKTQVIKLFSRFAT<br>SFKDYFKNRANCFSADDISSSSCHRIVNDNAEIFFSNALVYRRIVKSLSNDDIN<br>KISGDMKDSLKEMSLEEIYSYEKYGEFITQEGISFYNDICGKVNSFMNLYCQK<br>NKENKNLYKLQKLHKQILCIADTSYEVPYKFESDEEVYQSVNGFLDNISSKHI<br>VERLRKIGDNYNGYNLDKIYIVSKFYESVSQKTYRDWETINTALEIHYNNILP<br>GNGKSKADKVKKAVKNDLQKSITEINELVSNYKLCSDDNIKAETYIHEISHIL<br>NNFEAQELKYNPEIHLVESELKASELKNVLDVIMNAFHWCSVFMTEELVDK<br>DNNFYAELEEIYDEIYPVISLYNLVRNYVTQKPYSTKKIKLNFGIPTLADGWS<br>KSKEYSNNAIILMRDNLYYLGIFNAKNKPDKKIIEGNTSENKGDYKKMIYNL<br>LPGPNKMIPKVFLSSKTGVETYKPSAYILEGYKQNKHIKSSKDFDITFCHDLID<br>YFKNCIAIHPEWKNFGFDFSDTSTYEDISGFYREVELQGYKIDWTYISEKDIDL<br>LQEKGQLYLFQIYNKDFSKKSTGNDNLHTMYLKNLFSEENLKDIVLKLNGE<br>AEIFFRKSSIKNPIIHKKGSILVNRTYEAEEKDQFGNIQIVRKNIPENIYQELYK<br>YFNDKSDKELSDEAAKLKNVVGHHEAATNIVKDYRYTYDKYFLHMPITINF<br>KANKTGFINDRILQYIAKEKDLHVIGIDRGERNLIYVSVIDTCGNIVEQKSFNI<br>VNGYDYQIKLKQREGARQIARKEWKEIGKIKEIKEGYLSLVIHEISKMVIKYN<br>AIIVMEDLSYGFKKGRFKVERQVYQKFETMLINKLNYLVFKDISITENGGLLK<br>GYQLTYIPDKLKNVGHQCGCIFYVPAAYTSKIDPTTGFVNIFKFKDLTVDAK<br>REFIKKFDSIRYDSEKNLFCFTFDYNNFITQNTVMSKSSWSVYTYGVRIKRRF<br>VNGRFSNESDTIDITKDMEKTLEMTDINWRDGHDLRQDIIDYEIVQHIFEIFRL<br>TVQMRNSLSELEDRDYDRLISPVLNENNIFYDSAKAGDALPKDADANGAYCI<br>ALKGLYEIKQITENWKEDGKFSRDKLKISNKDWFDFIQNKRYL |

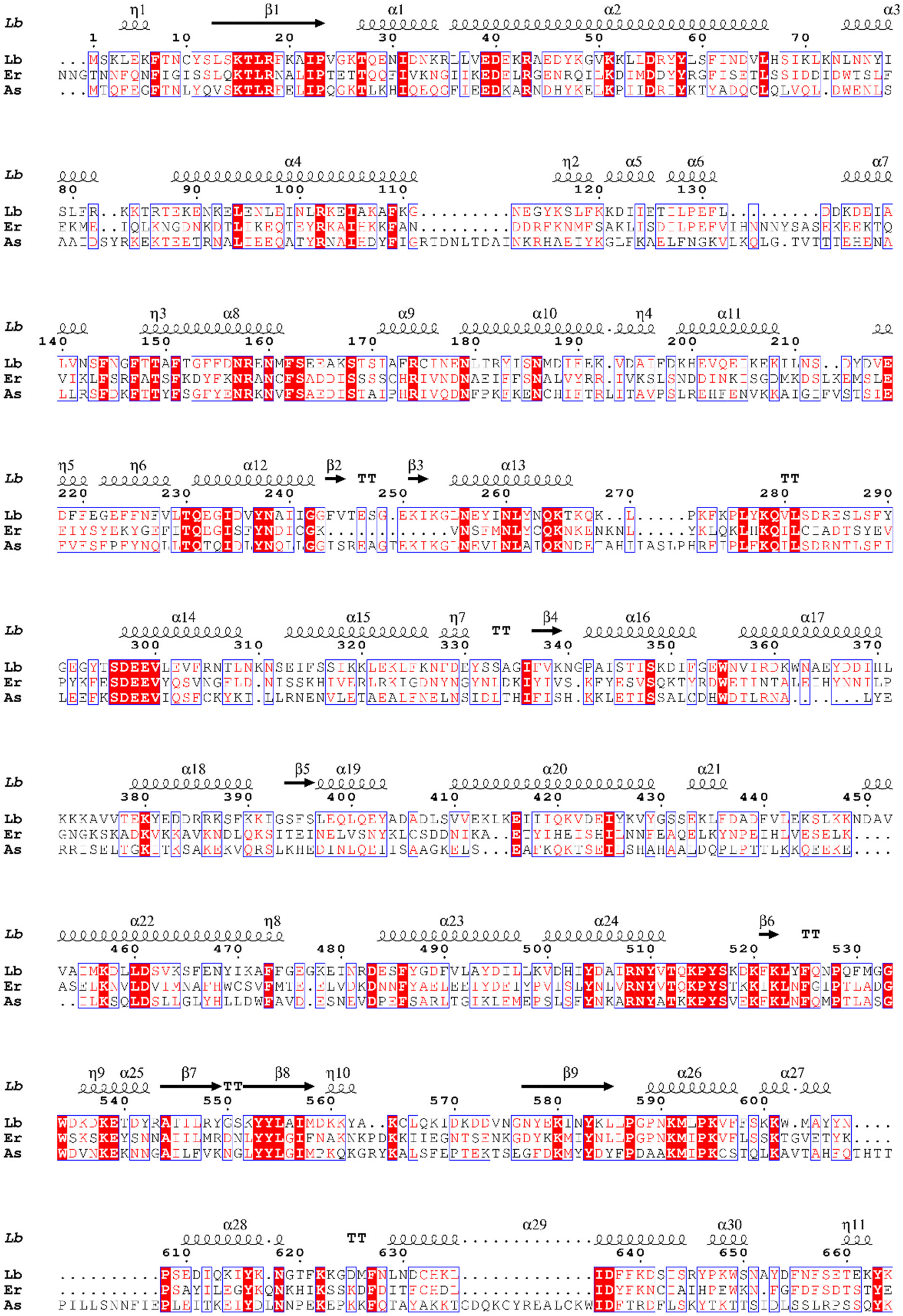
Lb
Er
As

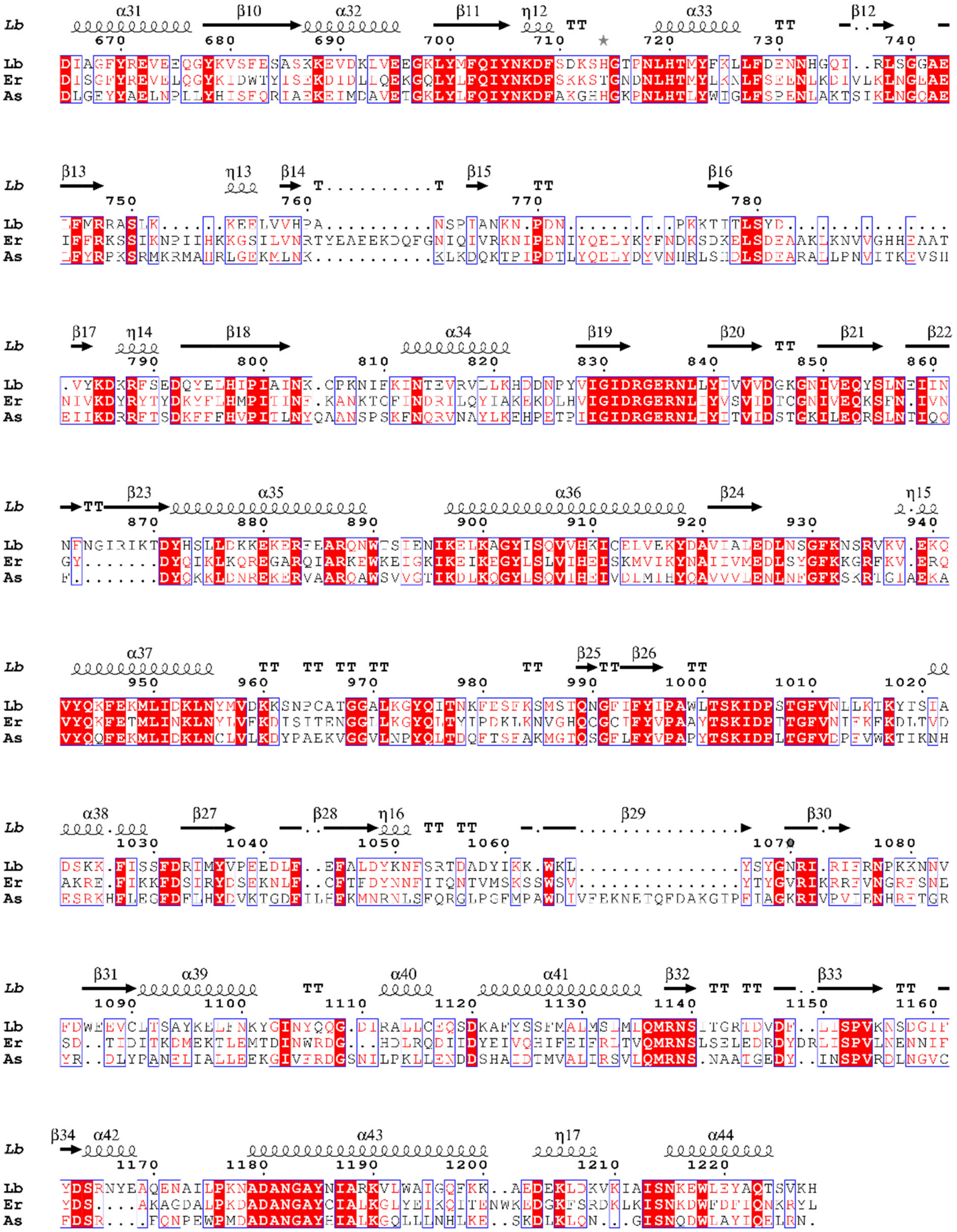

**Fig. S1:** Multiple sequence alignments of Lb, As, and Er Cas12a orthologs are shown. The alignment was performed using MultiAlin. The alignment file was imported into Esprit 3.0[48] and aligned against the LbCas12a structure (PDB: 5XUS)[49].

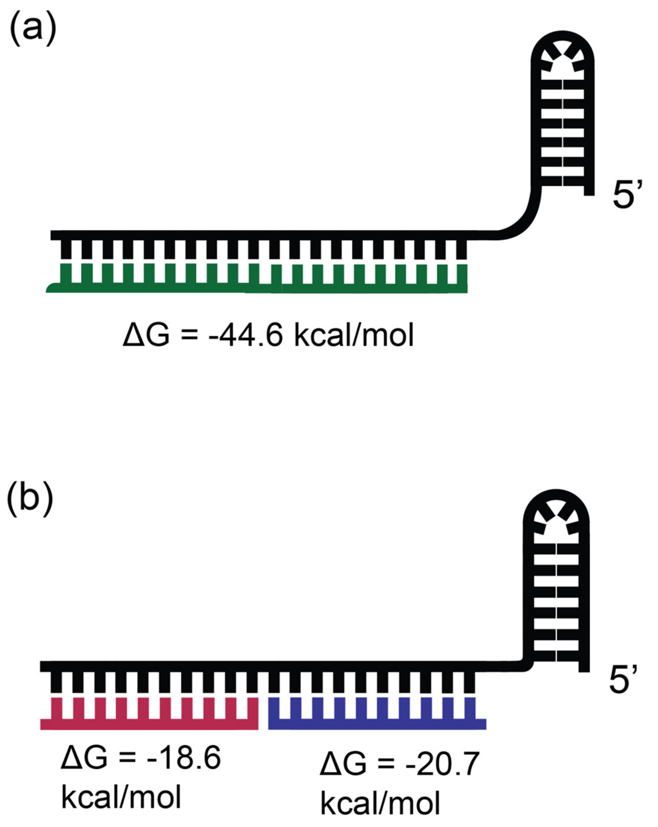


**Fig.S2:** Computationally predicted Gibbs free energy change for the binding of **(a)** a full-length 20-nt activator to the crRNA and **(b)** two short activators of length 10-nt each binding to different regions of the crRNA in a 'split-activator' fashion. Predictions were made using DINAMELT[50].

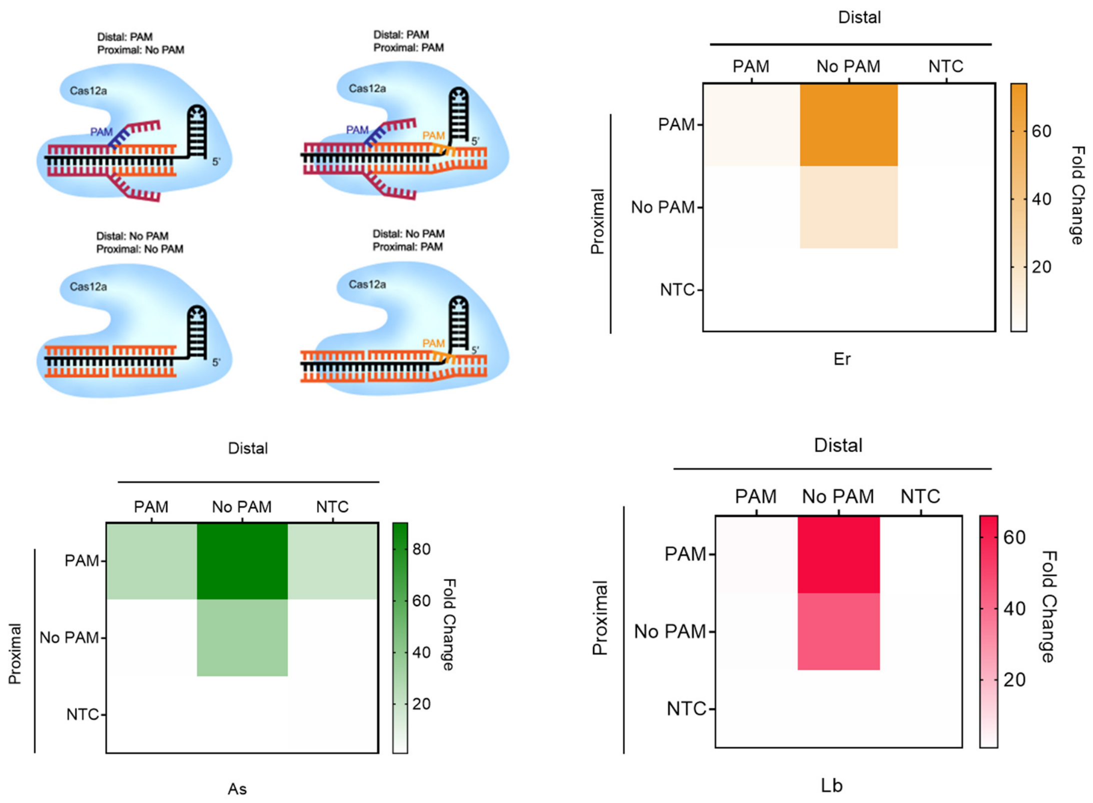


**Fig.S3:** *Trans*-cleavage activity of a combination of PAM- or no-PAM-containing double-stranded DNA activators binding at either the PAM-proximal (Pp) end or the PAM-distal (Pd) end of the crRNA in a combinatorial fashion. The heat map indicates the fold change in RFU compared to NTC at time t=60 min for Lb, As, and Er Cas12a orthologs (n=3).

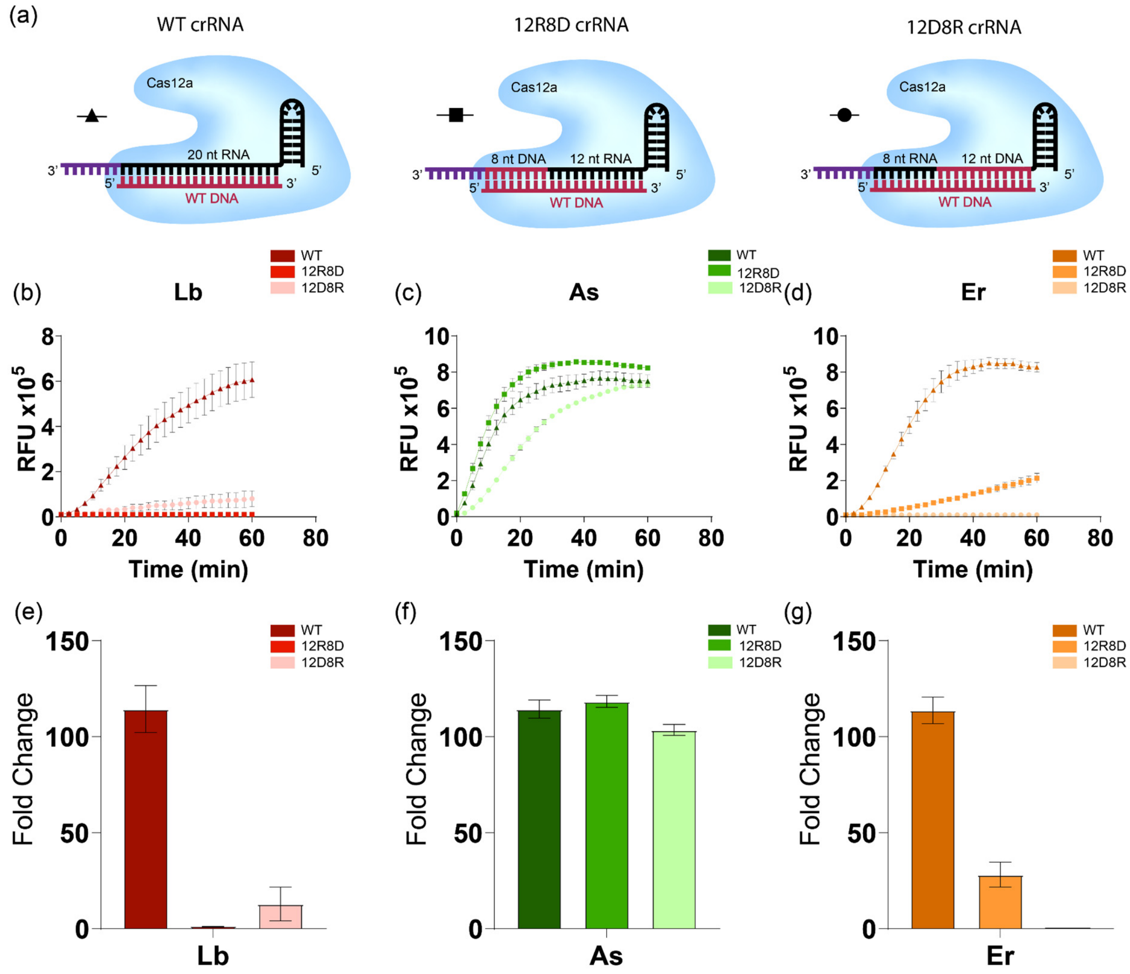


**Fig S4:** Chimeric DNA-RNA guides complexed with Cas12a. (a) Schematic representation of chimeric DNA-RNA hybrid crRNAs complexed with Cas12a and activated with WT ssDNA activators. Chimeric crRNA was designed by changing 12-nt near the PAM-proximal 5'-end of the crRNA to DNA (12D8R crRNA) and changing the PAM distal 8-nt end of the crRNA to DNA (12R8D crRNA). WT crRNA is represented in graphs WT crRNA is represented on graphs b-d by triangles, 12R8D crRNA is represented by squares, and 12D8R crRNA is represented by circles. (b-d) Relative RFU values of in vitro *trans*-cleavage assay with Cas12a orthologs (Lb- red, As- green, Er- orange) complexed with WT crRNA, 12D8R crRNA, and 12R8D crRNAs. (e-g) Fold change at 60 min is represented for each crRNA and three Cas proteins. The reactions contained 25 nM ssDNA GFP WT activator, 60 nM Cas12a, and 12 nM crRNA (WT, 12R8D, 12D8R). Reactions were incubated for 60 min at 37°C. Error bars represent SD (n=3).

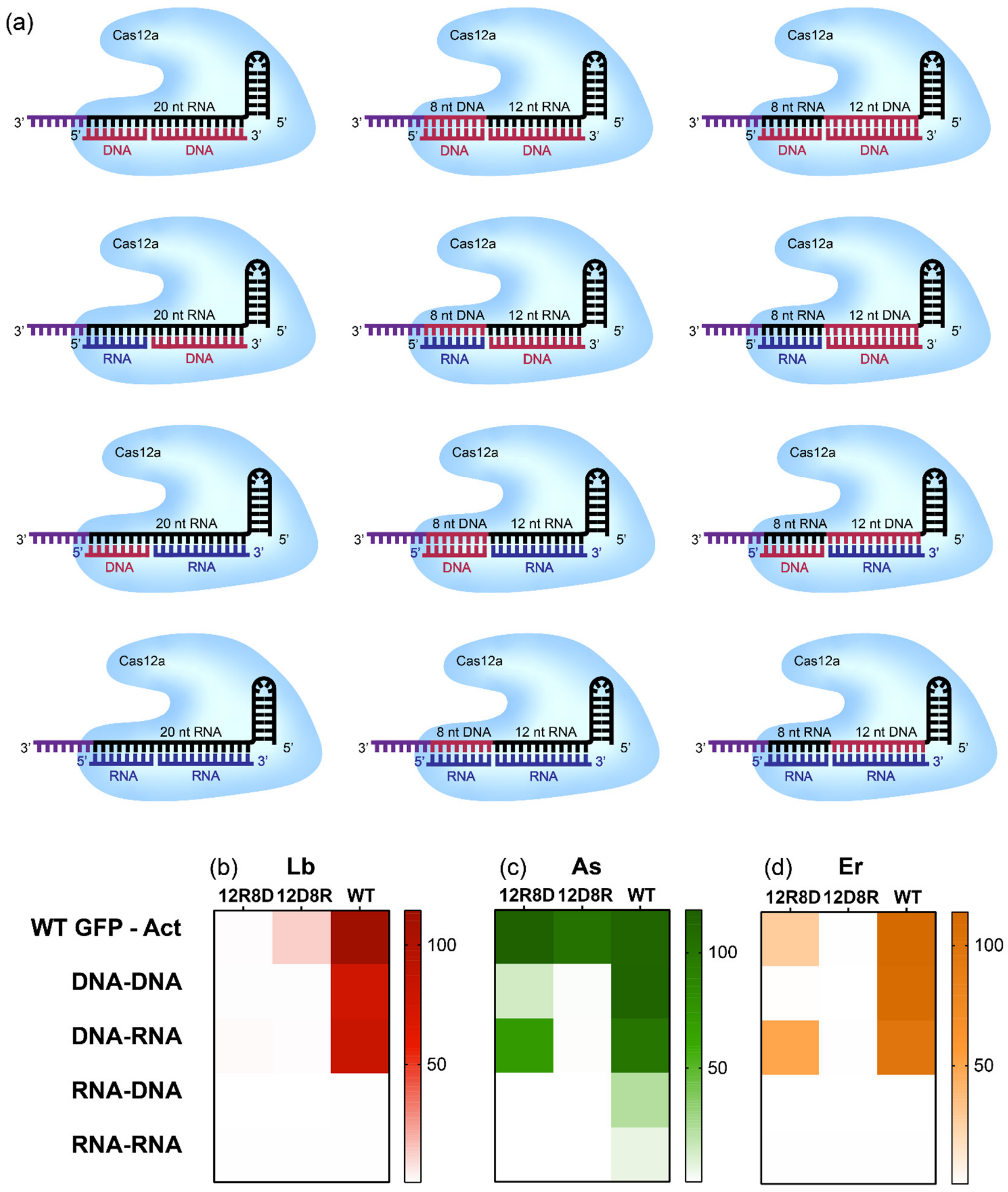


**Fig S5:** Reverse transcription-free RNA detection with Cas12a and 'split activator' mechanism. (a) Schematic representation of Cas12a complexed with chimeric crRNAs and activated by 'split activator' system. Chimeric crRNAs include 12D8R and 12R8D crRNAs as well as WT crRNA. Combinations of activators include ssDNA and RNA targeting the PAM proximal and distal locations on the crRNA. (b-d) Heat maps representing the fold changes of in vitro *trans*-cleavage assay with Cas12a orthologs complexed with WT and chimeric crRNAs. Combinatorial schemes for the 'split activator system' are seen in (a) (n=3).

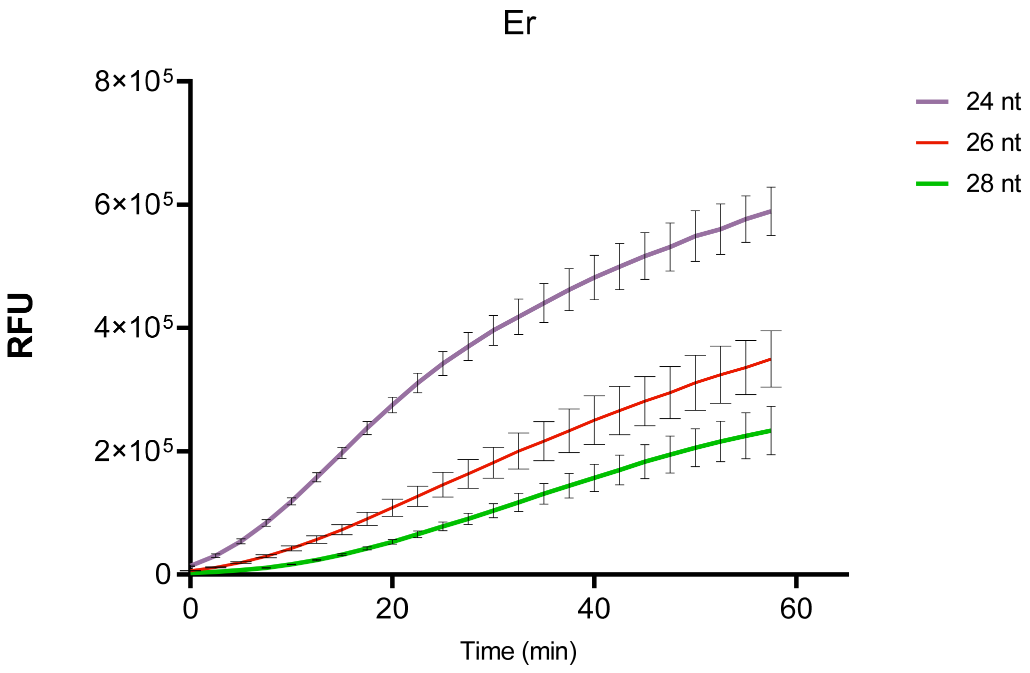


**Fig S6:** Increasing lengths of the crRNA ranging from 24-26 nt were tested for RNA detection with ErCas12a-based SAHARA. For each length of the crRNA, the S12 activator was kept at a constant length of 12-nt while the target RNA was varied from 12-16 nt to enable an increasing amount of target binding to the crRNA. The crRNA of length 24-nt that bound to 12-nt of S12 DNA and 12-nt of RNA target showed the highest activity.

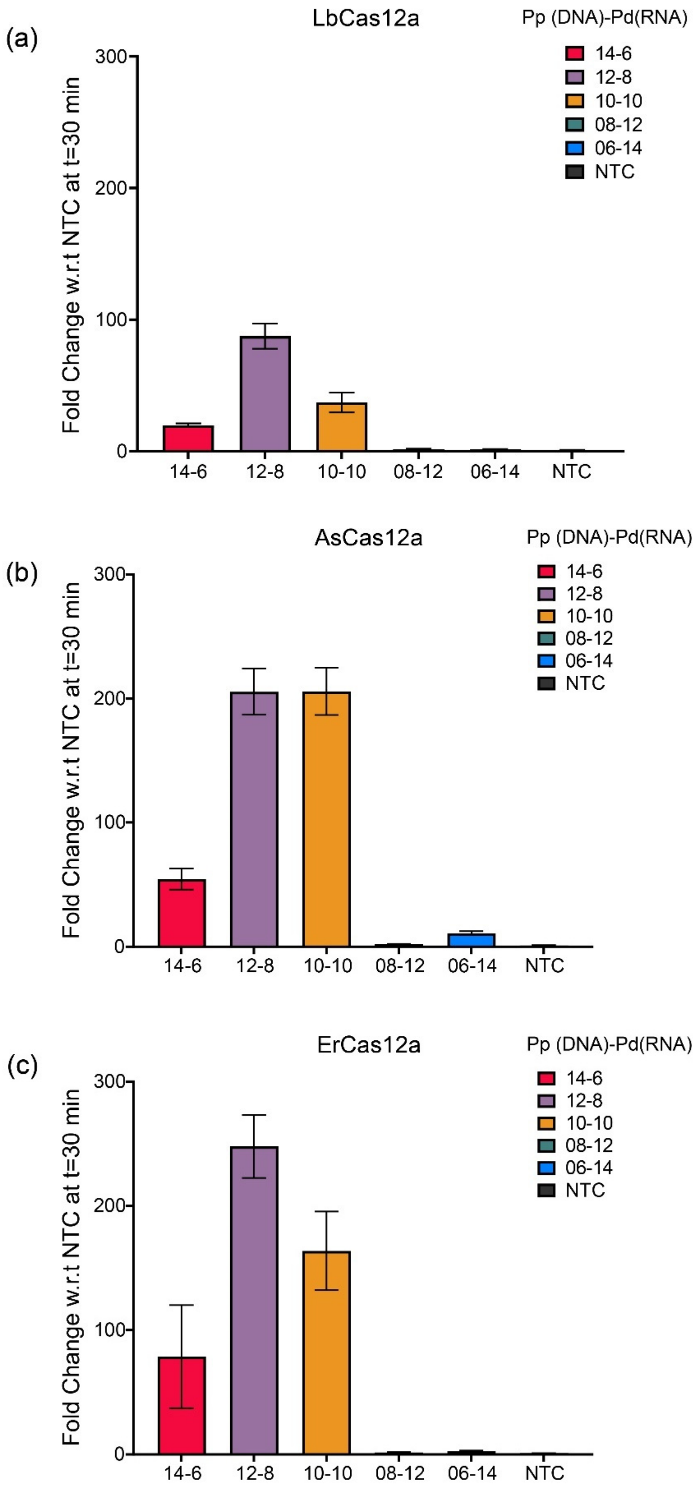


**Fig S7:** For a 20-nt crRNA, the length of the Pp binding DNA activator and the Pd binding RNA activator was varied from 6-14 nt. RNA detection was only tolerated for RNA activators of length 6nt-8nt, but not for RNA of length 12-nt or 14-nt.

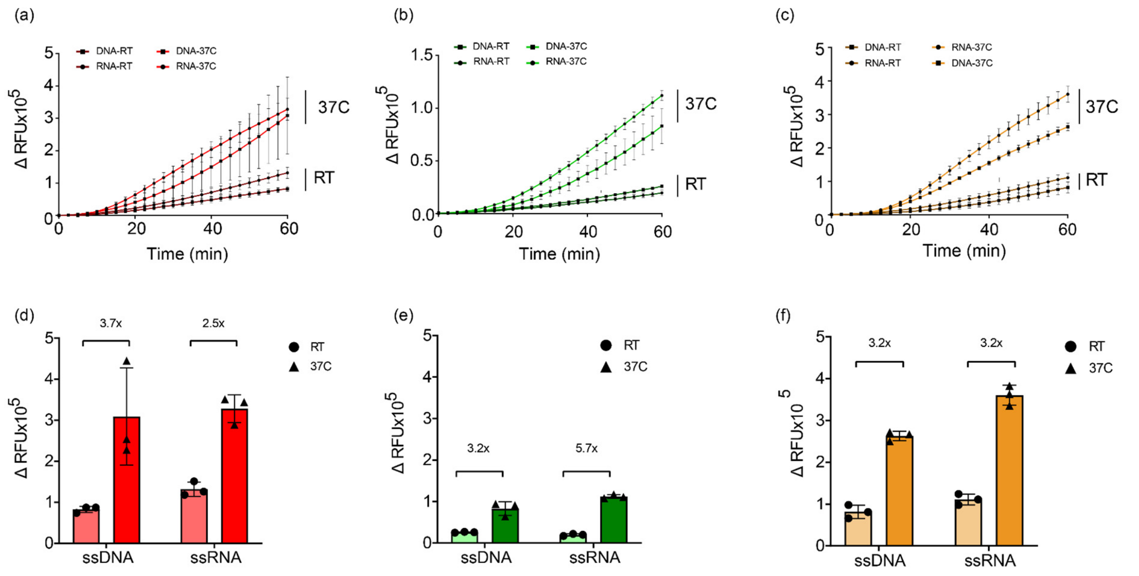


**Fig S8:** Detection of ssDNA or ssRNA sequences with SAHARA at different temperatures. **a-c:** Raw fluorescence data showing the *trans*-cleavage activity of SAHARA for the detection of an ssDNA or ssRNA sequence at either room temperature (RT) or 37°C for Lb (red), As (green), and Er (orange) orthologs. Error bars represent S.D. (n=3). **d-f:** Background subtracted raw fluorescence intensity for the detection of ssDNA or ssRNA sequences at RT or 37°C. Error bars represent S.D. (n=3).

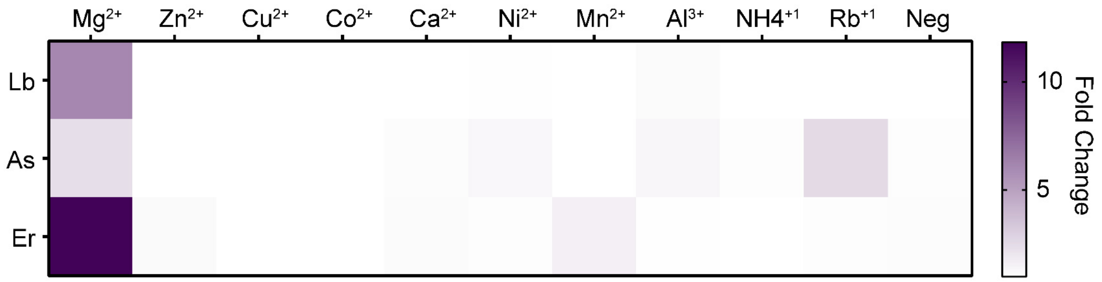


**Fig S9: Optimization of SAHARA with different divalent metal ions** (a) Effect of different metal ions on the SAHARA with Lb, As, and Er Cas12a enzymes. Negative control represents a no-salt buffer. Each metal ion buffer contains 3 mM of the respective metal salt. The heat map indicates fold change at time t=60 min (n=3).

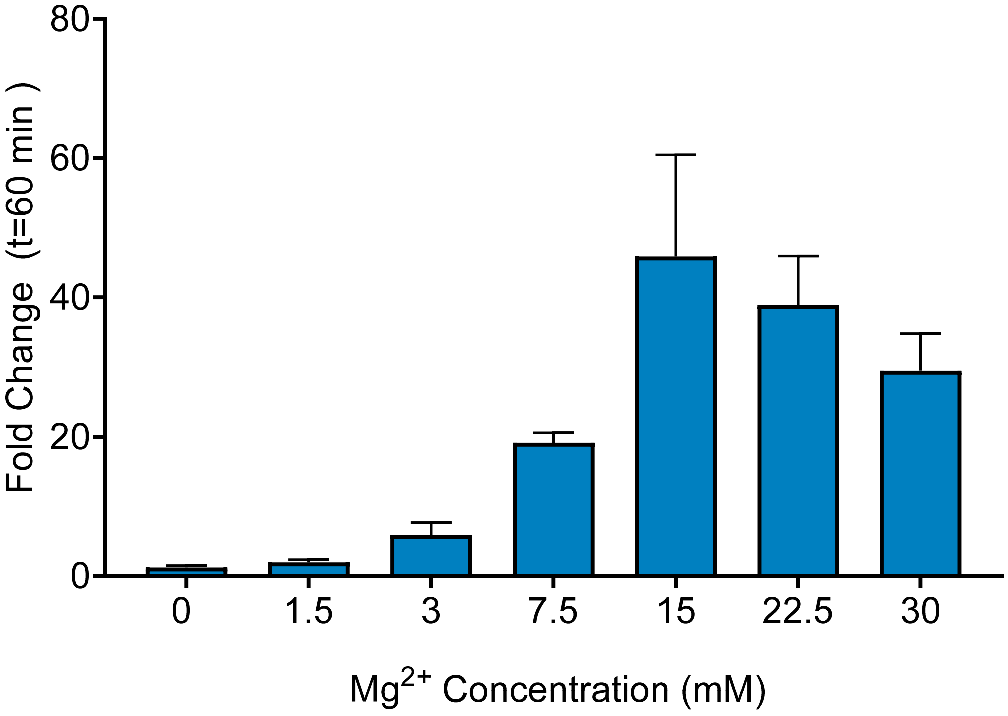


**Fig S10: Optimization of SAHARA with Mg ion concentration:** The *trans*-cleavage activity of SAHARA with ErCas12a under increasing $Mg^{2+}$ ion concentration is shown. The plot of fold change in RFU compared to NTC at t=60 min is shown. Error bars represent S.D. (n=3).

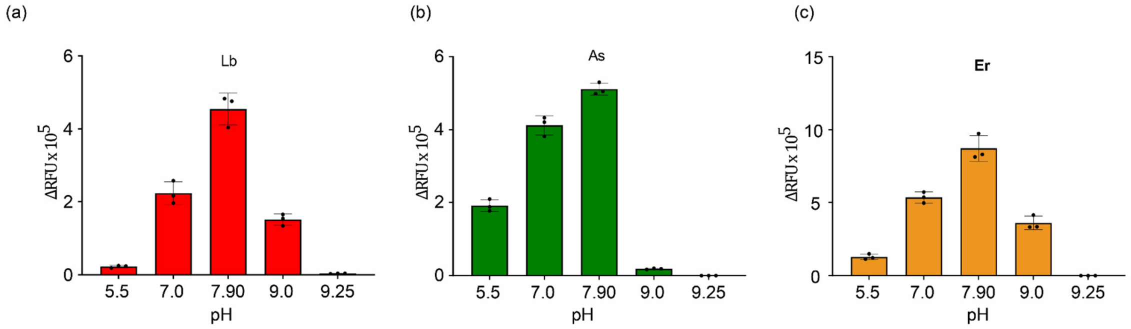


**Fig S11. Optimization of SAHARA with pH conditions a-c:** The effect of buffer pH on the *trans*-cleavage activity of SAHARA is shown. Bar graphs indicate background subtracted RFU at time t=60 min for Lb, As, and Er Cas12a orthologs at a different range of pH values. Error bars represent S.D. (n=3).

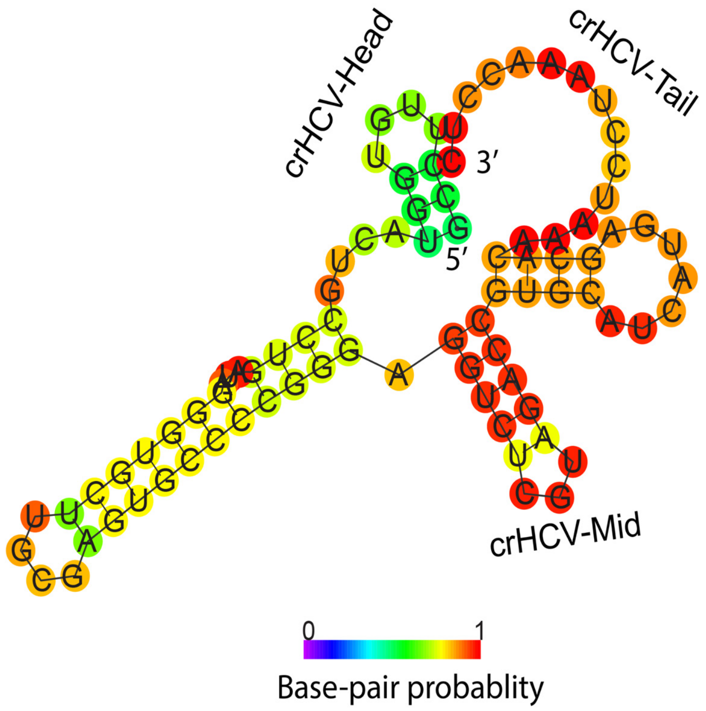


**Fig S12.** Computationally predicted secondary structure of the HCV polypeptide precursor targets. The three regions being targeted by SAHARA are labeled as HCV-Head (GCCUUGUGGUAC), HCV-Mid (AGACCGUGCAUC), and HCV-Tail (AUCCUAAACCUC) respectively. Predictions were done using the RNAFold webserver.

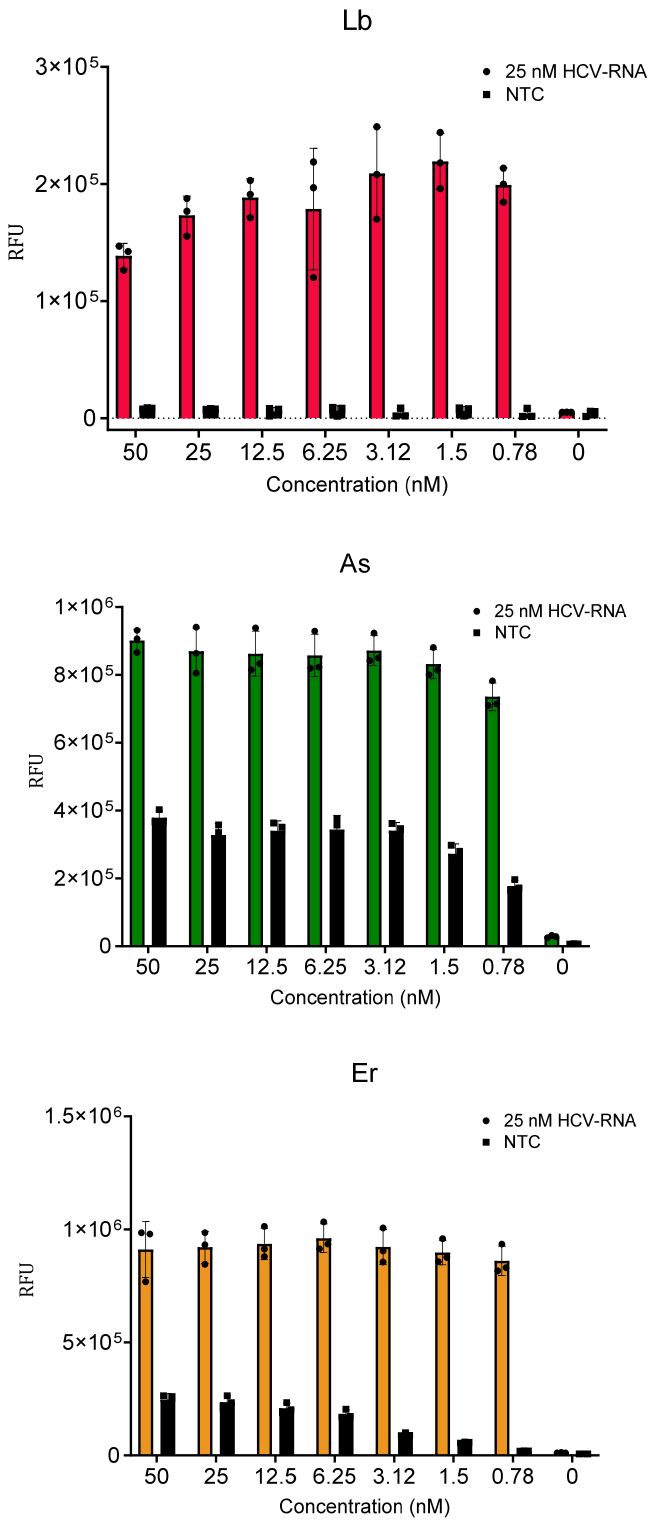


**Fig. S13:** Effect of S12 concentration ranging from 50 nM -780pM on the *trans*-cleavage activity of SAHARA is shown for Lb, As, and Er Cas12a orthologs. The plot of RFU at t=60 min in the presence or absence of 25 nM target HCV-RNA and different S12 concentrations is shown. Error bars represent S.D. (n=3).

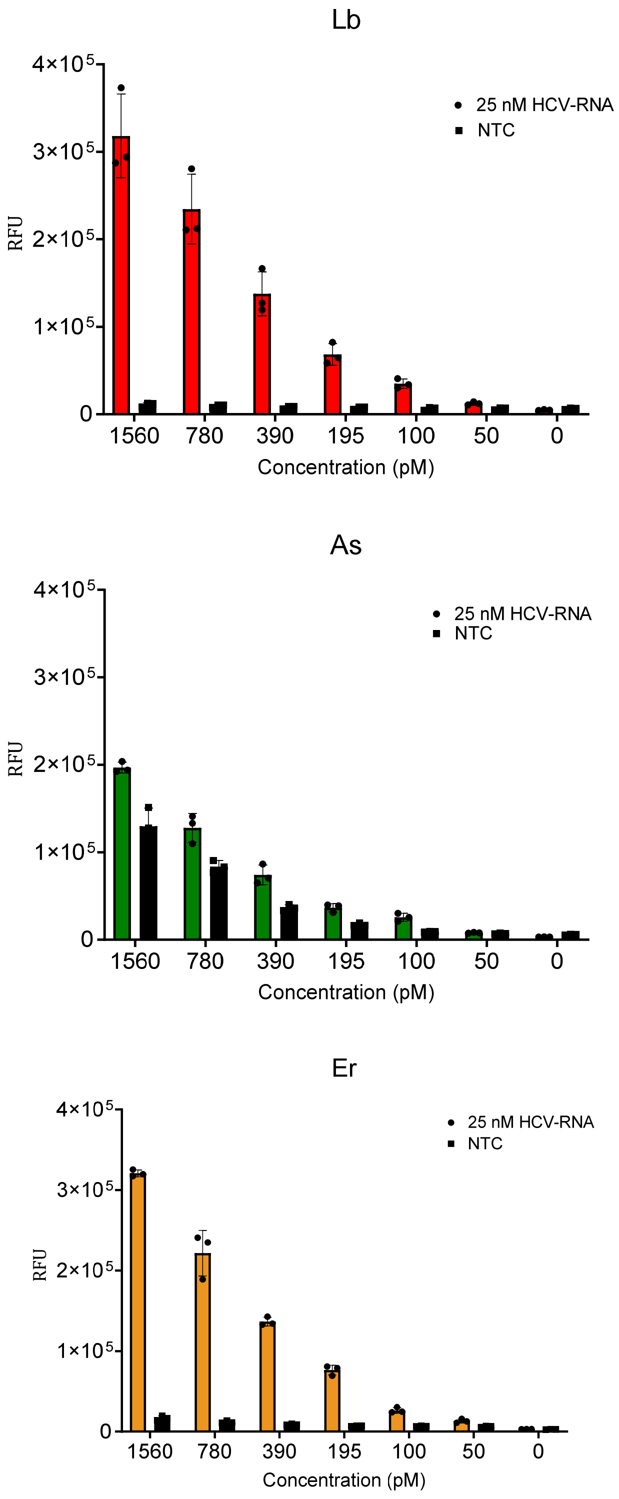


**Fig. S14:** Effect of S12 concentration ranging from 1.56 nM to 50 pM on the *trans*-cleavage activity of SAHARA is shown for Lb, As, and Er Cas12a orthologs. A plot of RFU at t=60 min in the presence or absence of 25 nM target HCV-RNA and different S12 concentrations is shown. Error bars represent S.D. (n=3).
